# Dynamic oligopeptide acquisition by the RagAB transporter from *Porphyromonas gingivalis*

**DOI:** 10.1101/755678

**Authors:** Mariusz Madej, Joshua B. R. White, Zuzanna Nowakowska, Shaun Rawson, Carsten Scavenius, Jan J. Enghild, Grzegorz P. Bereta, Karunakar Pothula, Ulrich Kleinekathoefer, Arnaud Baslé, Neil Ranson, Jan Potempa, Bert van den Berg

**Author notes:** The Harvard Cryo-Electron Microscopy Center for Structural Biology, Harvard Medical School, Boston, MA, USA. Authors contributed equally (Mariusz Madej, Joshua B. R. White).

## Abstract

*Porphyromonas gingivalis*, an asaccharolytic *Bacteroidetes*, is a keystone pathogen in human periodontitis that may also contribute to the development of other chronic inflammatory diseases, such as rheumatoid arthritis, cardiovascular disease and Alzheimer’s disease. *P. gingivalis* utilizes protease-generated peptides derived from extracellular proteins for growth, but how those peptides enter the cell is not clear. Here we identify RagAB as the outer membrane importer for peptides. X-ray crystal structures show that the transporter forms a dimeric RagA_2_B_2_ complex with the RagB substrate binding surface-anchored lipoprotein forming a closed lid on the TonB-dependent transporter RagA. Cryo-electron microscopy structures reveal the opening of the RagB lid and thus provide direct evidence for a “pedal bin” nutrient uptake mechanism. Together with mutagenesis, peptide binding studies and RagAB peptidomics, our work identifies RagAB as a dynamic OM oligopeptide acquisition machine with considerable substrate selectivity that is essential for the efficient utilisation of proteinaceous nutrients by *P. gingivalis*.

## Introduction

The Gram-negative Bacteroidetes are abundant members of the human microbiota, especially in the gut. Outside the gut, Bacteroidetes often cause disease, with the best-known examples being the oral Bacteroidetes *Porphyromonas gingivalis* and *Tannerella forsythia* that are part of the “red complex” involved in periodontitis^1^, the most prevalent infection-driven chronic inflammation in the Western world^2^. Accumulating evidence suggests a link between periodontitis and other chronic inflammatory diseases, including rheumatoid arthritis, Alzheimer’s disease, chronic obstructive pulmonary disease and cardiovascular disease^3–7^. Given this link, and the fastidious growth requirements of *P. gingivalis*, it is important to understand how this key pathogen thrives and causes dysbiosis of the oral microbiota in the biofilm on the tooth surface below the gum line, leading to inflammation and periodontal tissue destruction.

Unlike many human gut Bacteroidetes that specialise in degrading complex host and dietary glycans, *P. gingivalis* is asaccharolytic and exclusively utilises peptides for growth^8^. Those peptides are generated by multiple proteases, including several secreted endopeptidases and periplasmic di- and tri-peptidyl-peptidases^9,10^. The best-known *P. gingivalis* endopeptidases are the gingipains, large and abundant surface-anchored cysteine proteases with cumulative trypsin-like activity, which are essential for *P. gingivalis* virulence and growth on proteins as the sole source of carbon^11^. Crucially, it is not clear how gingipain-generated peptides are taken up by *P. gingivalis*. A role in this process has been proposed for the RagAB outer membrane (OM) protein complex^12^, which consists of a TonB dependent transporter (TBDT) RagA (PG_0185) and a substrate binding surface lipoprotein RagB (PG_0186). However, a recent crystal structure of *P. gingivalis* RagB was claimed to contain bound monosaccharides^13^, and a function related to polysaccharide utilisation was proposed based on sequence similarity with SusCD systems from gut Bacteroidetes^13–15^.

However, pairwise sequence identities between RagA/SusCs and RagB/SusDs are mostly very low (10-20%). Recent structural data for two SusCD complexes from *B. thetaiotaomicron* suggested that SusC and SusD form stable dimeric complexes (SusC_2_D_2_), with SusD capping the extracellular face of the SusC transporter. Molecular dynamics (MD) simulations and electrophysiology studies led to the proposal that nutrient uptake likely occurs via a hinge-like opening of the SusD lid, a so-called pedal bin mechanism^16^. However, the molecular details of nutrient acquisition by SusCD-like complexes remain unclear, as do any differences between putative OM peptide and glycan transporters.

Here we report comprehensive structural and functional studies of RagAB purified from *P. gingivalis* W83 via X-ray crystallography and single particle cryo-electron microscopy (cryo-EM). The crystal structure shows a dimeric RagA_2_B_2_ complex in which RagB, in a manner reminiscent to SusD-like proteins, caps the RagA transporter. This closed conformation generates a large internal chamber that is occupied by co-purified bound peptides at the RagAB interface. Remarkably, and in sharp contrast to the crystal structure of RagAB (and indeed those of SusCDs), cryo-EM reveals the large conformational changes that can occur, with three distinct states of the RagA_2_B_2_ transporter, *i.e*. open-open, open-closed and closed-closed, present in a single dataset of the detergent-solubilized complex. Together with structure-based site-directed mutagenesis analyses, peptide binding studies and RagAB peptidomics, we show that RagAB is a highly dynamic OM oligopeptide acquisition machine, with considerable substrate selectivity that is essential for the utilisation of protein substrates by *P. gingivalis*.

## Results

### Purification and X-ray crystal structure determination of RagAB from *P. gingivalis*

Given that SusC proteins from *Bacteroides* spp. do not express in the *E. coli* OM^16^, we purified RagAB directly from *P. gingivalis* W83 KRAB (Δ*kgp*/Δ*rgpA*/Δ*rgpB*). This strain lacks gingipains, reducing proteolysis of many OM proteins. RagAB is together with the OmpA-like Omp40-41 complex the most abundant OM protein in *P. gingivalis*. With the exception of the 230 kDa Hemagglutinin A (HagA) that is abundant as the full-length protein in the absence of gingipains, no co-purifying proteins are present (Fig. 1a). This indicates that the RagAB complex represents the complete transporter (indeed the operon contains only *ragA* and *ragB*). Diffracting crystals were obtained by vapour diffusion, and the structure was solved by molecular replacement using data to 3.4 Å resolution (Methods and Supplementary Table 1).

**Figure 1.**
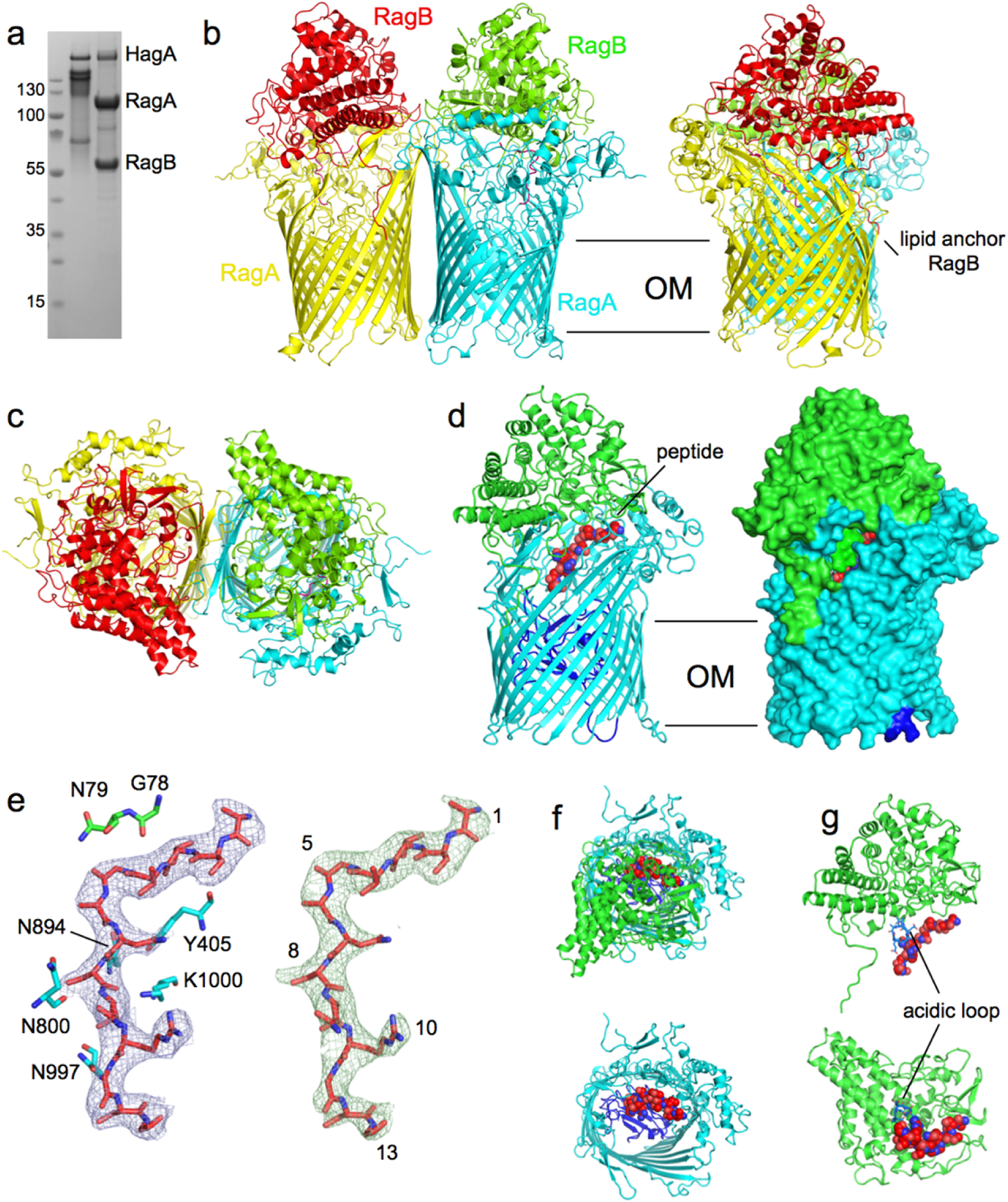
Crystal structure of the RagA_2_B_2_ transporter shows bound peptides. **a**, SDS-PAGE gel showing purified RagAB from W83 KRAB before (left lane) and after boiling in SDS-PAGE sample buffer. **b,c,** Views of RagA_2_B_2_ from the plane of the OM (**b**) and from the extracellular space (**c**). **d**, Side views of RagAB (right panel; surface representation) showing the bound peptide as a red space filling model. The plug domain of the RagA TBDT (cyan) is dark blue. **e**, 2Fo-Fc density (blue mesh; 1.0 σ, carve = 1.8) of the peptide after final refinement. Protein residues that likely form hydrogen bonds with the peptide backbone are shown. The right panel shows Fo-Fc density (green mesh; 3.0 σ, carve = 1.8) obtained after removal of the peptide from the model. Arbitrary peptide residue numbering is indicated. **f**, Extracellular views of RagAB with (top panel) and without the RagB lid (green). **g**, Side view (top panel) and view from the direction of RagA for RagB and the bound peptide. The acidic loop of RagB is coloured blue. Structural figures were made with Pymol^17^. Several protein residues likely form hydrogen bonds with the backbone of the peptide substrate.

The structure shows that RagAB is dimeric (RagA_2_B_2_), with the same subunit arrangement and architecture (Fig. 1) observed previously for two SusCD complexes from *Bacteroides thetaiotaomicron*^16^. RagB caps RagA, burying an extensive surface area (~ 3800 Å^2^) and forming a closed complex with a large internal cavity. Inspection of this cavity reveals clearly resolved density for an elongated, ~40 Å long molecule that is bound at the RagAB interface. The similarity with the neighbouring protein electron density is striking, and strongly suggests that the bound molecule is a peptide of ~13 residues in length (Figs. 1d-g), although the densities for most side chains are truncated after the Cβ or Cγ positions. Exceptions are residues 9 and 10 of the peptide, where well-defined sidechain density comes within hydrogen bonding distance of the side chains of residues D99 and D101 in RagB. We modelled the peptide as A^1^STTG^5^ANSQR^10^GSG^13^ (Fig. 1e). D99 and D101 are part of an acidic loop insertion (Asp^99^-Glu-Asp-Glu^102^) into an α-helix in the ligand binding site of RagB that protrudes into the RagAB binding cavity (Fig. 1g). From the acidic nature of this loop we speculate that RagAB of *P. gingivalis* W83 may prefer basic peptides for uptake. Interestingly, the acidic loop is absent in a number of RagB orthologs, including that from strain ATCC 33277 (Supplementary Fig. 1). The backbone of the peptide has an occupancy that is only slightly lower than neighbouring RagA and RagB chains, suggesting that this density most likely corresponds to an ensemble of peptides with different sequences, but similar backbone conformations, hereafter referred to as “the peptide”.

These are the carbonyl of G78 and the side chain of N79 from RagB, which interact with peptide residues 4 and 5 respectively (Fig. 1e). From RagA, the carbonyls of Y405 and N800 form hydrogen bonds with peptide residues 3 and 9. The side chain of N894 forms hydrogen bonds with both residues 6 and 8, and may therefore be particularly important for substrate binding. Finally, the side chains of N997 and K1000 likely interact with peptide residues 11 and 8 respectively (Fig. 1e). Thus, RagAB residues interact with the backbone of 8 out of the 13 visible peptide residues, and most of those interactions are provided by RagA. Indeed, a PISA interface analysis^18^ shows that 26 RagA residues form an interface with the peptide compared to only 8 for RagB, generating interface areas of 620 and 240 Å^2^ with RagA and RagB respectively. The PISA CSS (complexation significance score) is 1.0 for peptide-RagA while it is only 0.014 for peptide-RagB. This suggests that the observed co-crystal structure represents a state where the ligand has been partially transferred from an initial, presumably low(er)-affinity binding site on RagB to a high(er)-affinity binding site in the RagAB complex, allowing co-purification.

### Conformational changes in RagAB

The crystal structure of RagAB, and those previously determined for SusCD complexes, are in very similar states in which both TBDT barrels are closed on the extracellular side by their RagB (or SusD) caps, even when no substrate is bound^16^. This suggests either that the closed state is energetically favourable, or that crystallisation selects for closed states from a conformational ensemble in solution. We therefore investigated the structure of detergent-solubilized RagAB in solution using single particle cryo-EM (Methods). Following initial 2D classification, it was apparent that RagAB is also a dimer in solution, thus demonstrating that the dimeric complexes observed via X-ray crystallography are not artefactual. Strikingly, it was also immediately apparent that multiple conformations of the RagA_2_B_2_ dimer were present in the data, and after further classification steps, three distinct conformations were identified. Following 3D reconstruction and refinement, three structures were obtained at near-atomic resolution, corresponding to the three possible combinations of open and closed dimeric transporters: closed-closed (CC; 3.3 Å), open-closed (OC; 3.3 Å) and open-open (OO; 3.4 Å) (Figs. 2 and 3; Supplementary Table 2).

**Figure 2.**
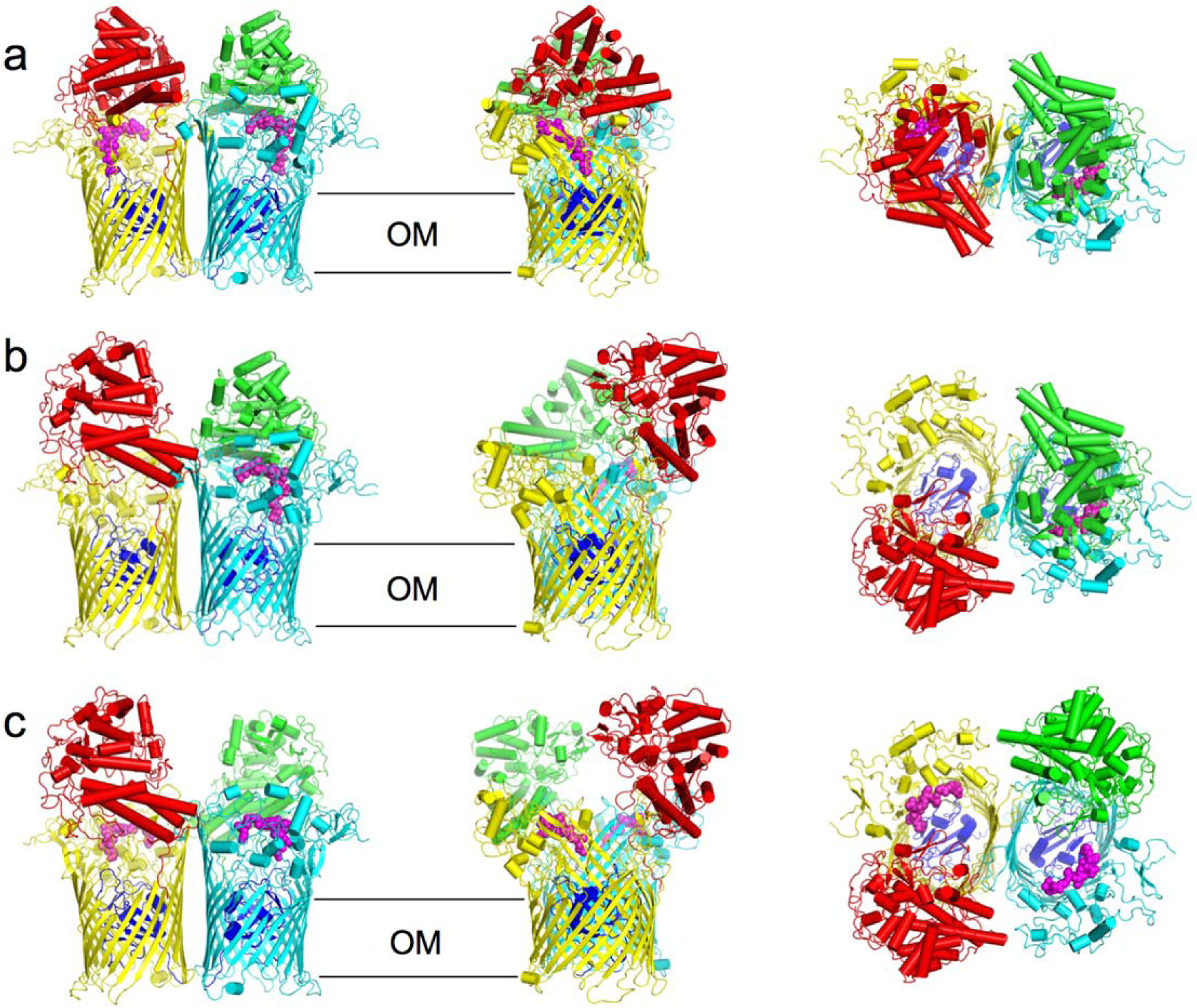
Different conformations of the RagA_2_B_2_ transporter revealed by cryo-EM. **a-c**, Cartoon views for closed-closed RagA_2_B_2_ (CC; **a**), open-closed (OC; **b**) and open-open (OO; **c**). Views are from the plane of the OM (left and middle panels) and from the extracellular space (right panel). Bound peptide is shown as space filling models in magenta. The plug domains of RagA are coloured dark blue.

**Figure 3.**
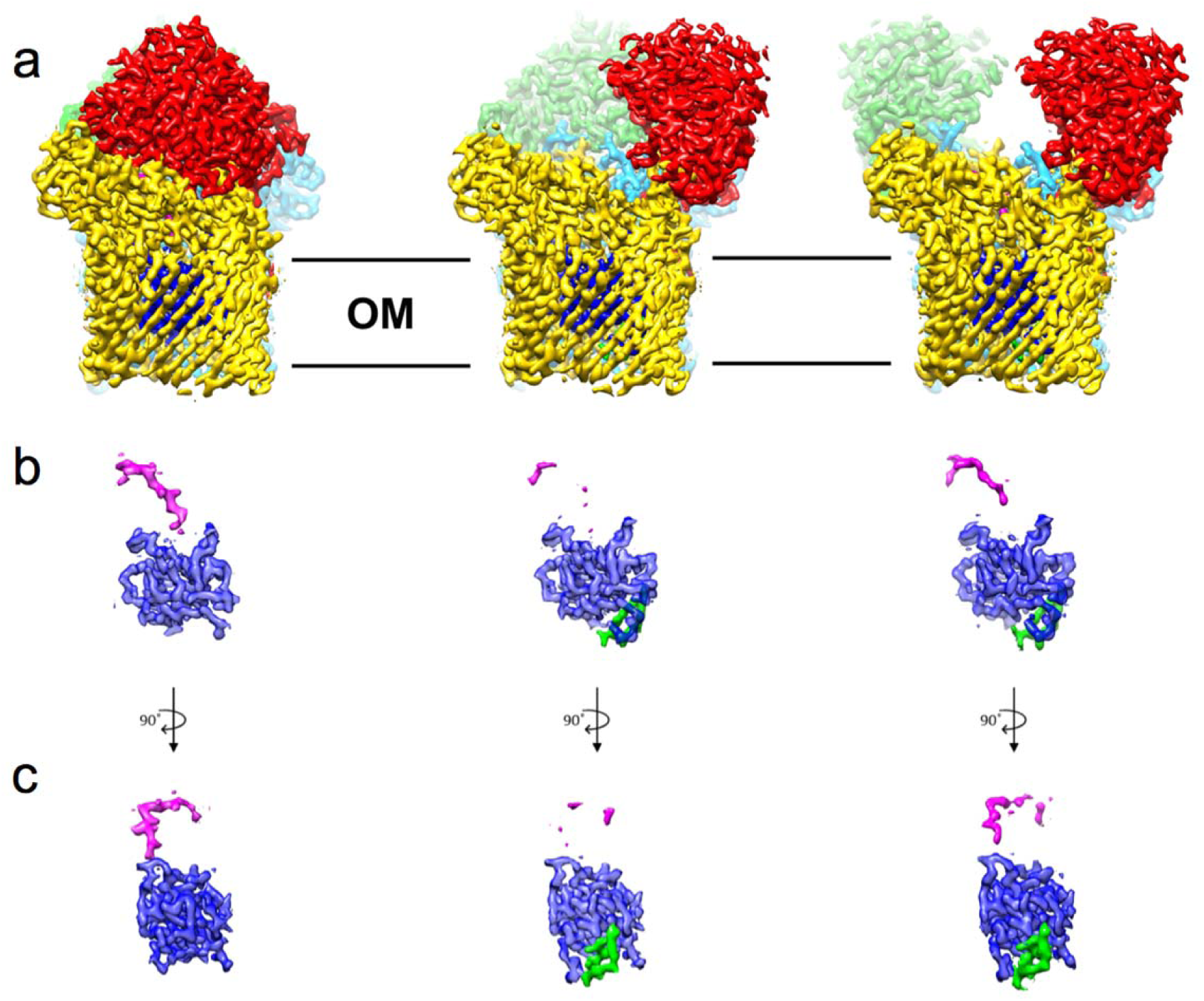
Plug and peptide dynamics in RagA_2_B_2_ revealed via cryo-EM. **a**, Density maps viewed from the membrane plane of the closed-closed, open-closed and open-open RagA_2_B_2_ complexes coloured as in Figure 2. **b,c**, Isolated density for the plug domain and the bound peptide ensemble viewed as in (**a**) and after a 90 degree clockwise rotation (**c)**, Additional density for the plug domains of open RagAB complexes is shown in green.

The CC state is essentially identical to the X-ray crystal structure (backbone r.m.s.d. values of ~0.6 Å), and the bound peptide occupies the same position. In the OO state, each RagB lid has undergone a very substantial conformational rearrangement that swings RagB upward, exposing the peptide binding site and the plug domain within the barrel interior. The bound peptide is present in both barrels of the OO state, at the same position as in the CC state, but with lower occupancy (Fig. 3). This is consistent with the PISA interface analysis showing that the observed peptide binding site is mainly formed by RagA.

The most interesting state is the open-closed transporter (OC), in which the internal symmetry of the RagA_2_B_2_ complex has been broken. The density for bound peptide in the open barrel of the OC RagAB is much weaker compared to those in either barrel of the OO state (Fig. 3), even when both maps are generated by assuming C1 symmetry. The OC state unambiguously shows that the RagB lids in the dimeric complex can open and close separately, *i.e*. the “pedal bins” can function independently. The open and closed transporters of the OC state are identical to those in the CC and OO states, respectively. Fascinatingly, there is no evidence for “intermediate” states in the opening of the transporter: essentially all of the individual RagAB pairs imaged are either open or closed, and these are dimerised to give each of the three possible states observed (CC, OC and OO). This suggests that either the dynamics of lid opening and closing are very rapid, and/or that the energy landscape for opening and closing is very rugged, with distinct stable minima only existing for the three discrete conformational states seen in the EM experiment. MD simulations performed prior to obtaining the EM structures show a similar, albeit less wide opening of the RagB lid upon removal of the peptide (Supplementary Fig. 2).

### Lid opening involves a pivot around the N-terminus of RagB

Superposition of the RagA subunits of the open and closed states reveals that the N-terminal ~10 residues of the RagB lid and the lipid anchor at the back of the complex remain stationary during lid opening (Fig. 4a), acting as a hinge point about which the rest of the protein moves as a rigid body. Lid opening is a complex, 3D movement involving a rotation both about the hinge point and around an axis through the RagB subunit (Supplementary Movie 1). This results in displacements of up to 45 Å for main chain atoms at the front of the complex, furthest away from the RagB N-terminus (Figs. 4 a-c). For RagA, the conformational changes upon lid movement are mostly confined to those parts of the protein that continue to interact with RagB at the back of the complex, and consist of movements of up to 13-14 Å for main chain atoms in extracellular loops L7, L8 and L9 (Fig. 4d).

**Figure 4.**
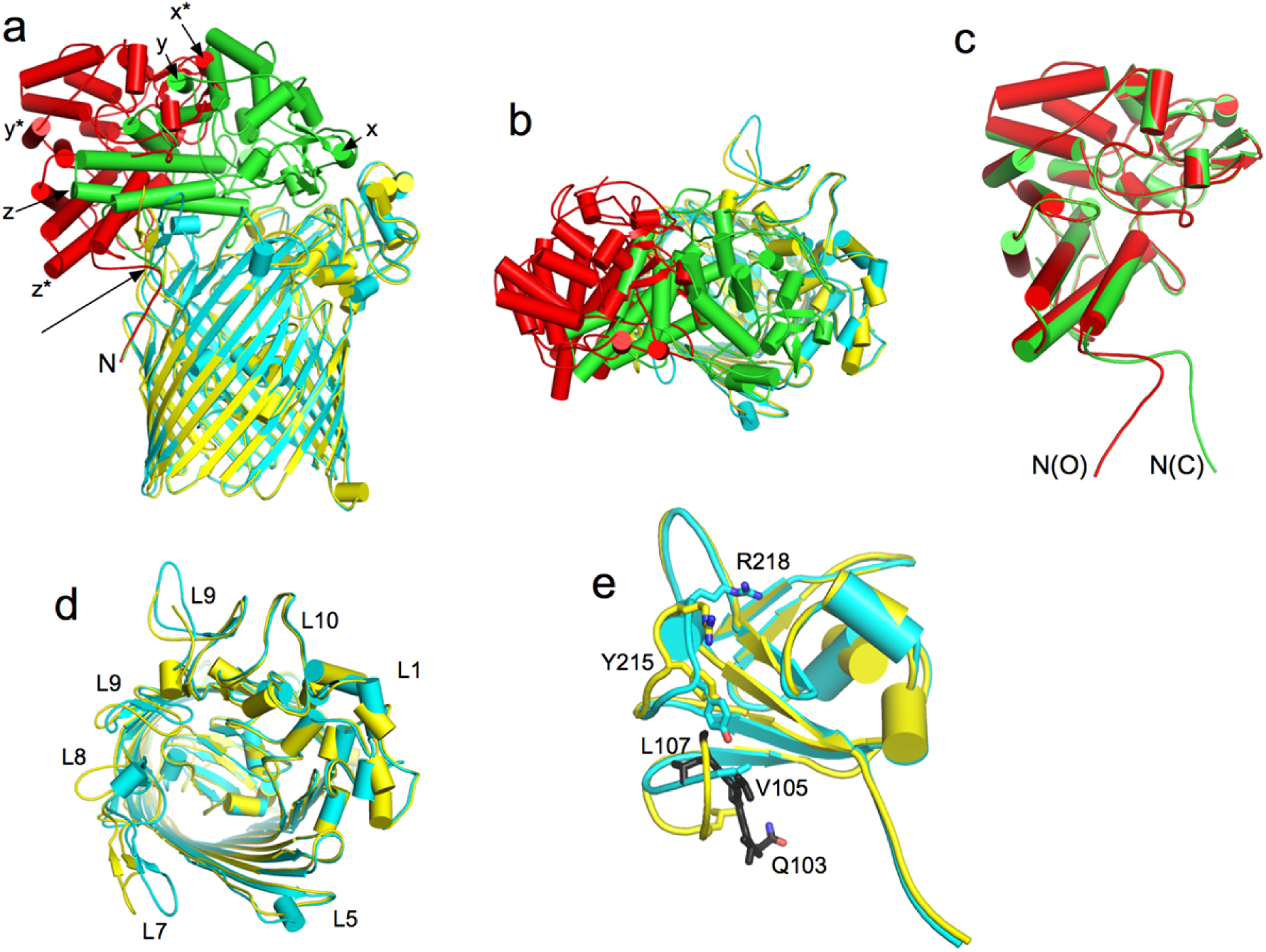
RagB moves as a rigid body during lid opening. **a**, Superposition of RagA subunits in the open (yellow) and closed (cyan) RagAB complexes, showing the rigid-body movement of RagB. Equivalent points are indicated by x,y,z (closed RagB; green) and by x*, y*, z* (open RagB; red). The arrow indicates the approximate pivot point in the N-terminus of RagB at the back of the complex. **b**, Superposition as in (**a**), viewed from the extracellular side. **c**, Superposition of the open and closed states of RagB, with the N-termini indicated. **d**, Extracellular view of superposed RagA, with selected loops labelled. **e**, Side view comparison of the RagA plug domains. The periplasmic space is at the bottom of the panel. Selected residues that show substantial conformational changes are shown as stick models. The TonB box of open RagAB (yellow) is coloured black and residues are shown as sticks. The TonB box is invisible in the closed state (cyan).

### The open-closed complex reveals changes in the RagA plug domain

In the consensus model of TonB-dependent transport^14^, extracellular substrate binding generates conformational changes in the plug domain of the TBDT that are transmitted to its N-terminal Ton box. This increases accessibility of the Ton box from the periplasm, allowing interaction with TonB in the inner membrane. This ensures that only substrate-loaded transporters are TonB substrates, avoiding futile transport cycles with empty transporters. In crystal structures of TBDTs, substrate binding is generally manifested by the Ton box being visible in the structures of “apo” TBDTs due to interactions with other parts of the plug, and therefore not being available for TonB binding. In structures of ligand-bound TBDTs these interactions are lost, leading to increased mobility of the N-terminus and loss of Ton box electron density. Our cryo-EM structure of the OC state offers an unbiased structural view that relates the occupancy of the ligand binding site to changes in the plug domain (Figs. 3 and 4e). In the open side, the first visible residue is Q103 at the start of the TonB box, whereas in the closed side (and in the crystal and other EM structures), the density starts at L115. The conformation of L115-S119 is also different from that in the open complex with shifts for L115 as large as 10 Å. Interestingly, the plug region A211-A219 is also different in the two states, and especially the pronounced shift for R218 (~9 Å for the head group; Fig. 4e) may be important given its location at the bottom of the binding cavity. A211-A219 contacts the region following the Ton box, highlighting a potential allosteric route via binding site occupancy could affect the conformation and dynamics of the TonB box. Such a mechanism is also consistent with previous AFM data suggesting that the plugs of TBDTs consist of two domains^19^ : an N-terminal, force-labile domain that is removed by TonB to form a channel and a more stable C-terminal domain that would include R218 in RagA and which would signal occupancy of the binding site to the TonB box. The difference in dynamics of the N-terminus of RagA is very evident in unsharpened maps of the OC state contoured at low levels, which clearly show globular density connected to the plug domain only in the *open* state (Supplementary Fig. 3). This density corresponds to the ~80-residue N-terminal extension of unknown function (DUF4480) that precedes the TonB box in RagA and other Bacteroidetes TBDTs. In the closed state, the DUF and Ton box are likely too mobile to be detected in the cryo-EM density maps.

### RagAB is important for growth of *P. gingivalis* on proteins as carbon source

We initially assessed the ability of several *P. gingivalis* strains to grow on minimal medium supplemented with BSA as a sole carbon source (BSA-MM). Interestingly, while growth on rich medium is identical, robust growth on BSA-MM is observed only for the *rag-1* locus strains W83 and A7436^20^. By contrast, *rag-4* strains ATCC33277, HG66 and 381 failed to grow on BSA (Fig. 5a). The RagAB complexes of strains that grow on BSA (*rag-1*) have the RagB acidic loop (Fig. 1 and Supplementary Fig. 1), whereas this insertion is lacking in *rag-4* strains. This suggests that relatively small structural differences in RagAB could alter substrate specificity sufficiently to prevent growth on the relatively small set of peptides that can be generated from BSA. To investigate this and define the role of RagAB in peptide utilisation we constructed clean deletions in *P. gingivalis* W83 for *ragA*, *ragB*, and the complete *ragAB* locus and grew the resulting strains on BSA-MM.

**Figure 5.**
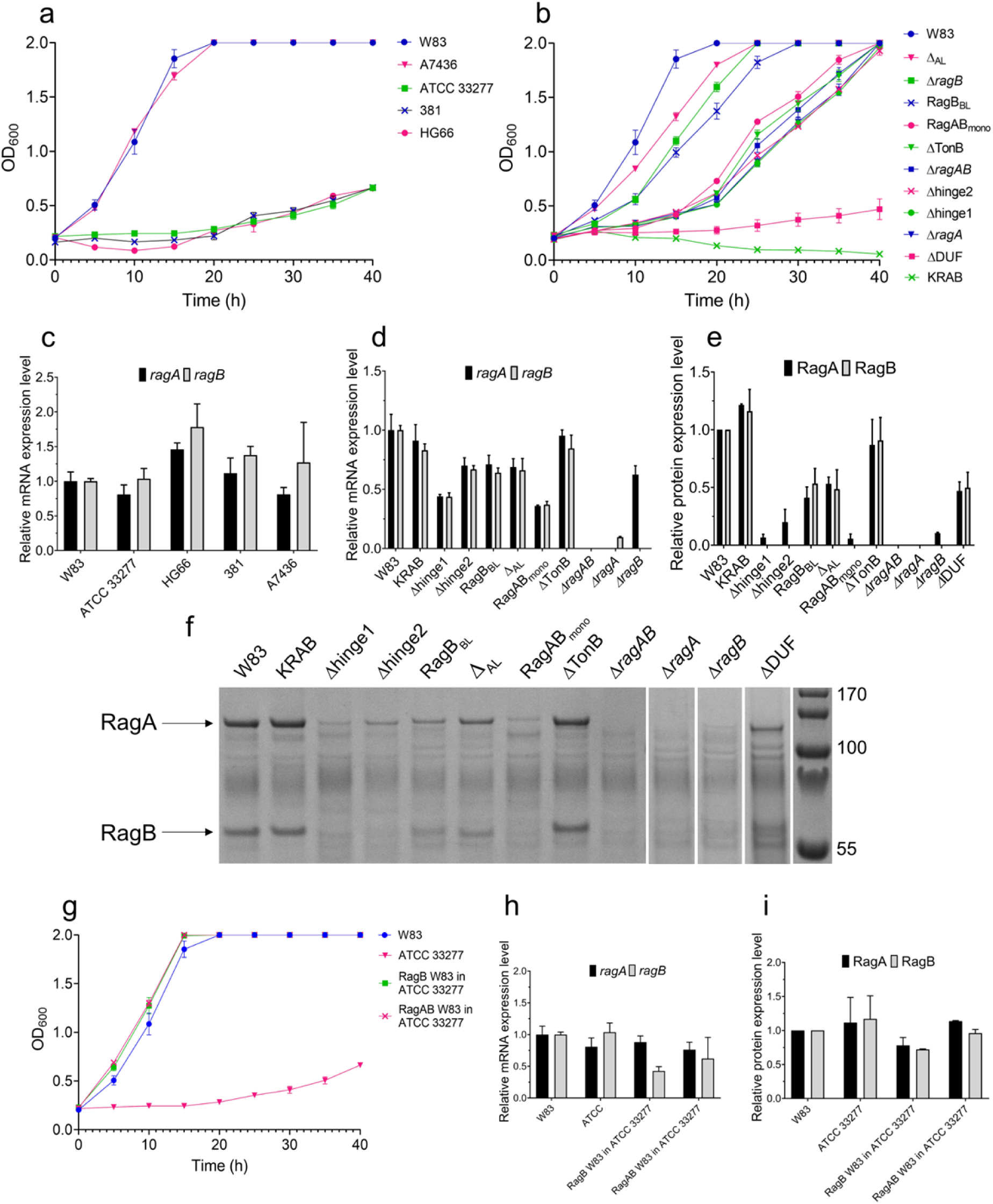
RagAB is required for growth on BSA. **a**, Representative growth curves (n = 3, mean ± standard deviation) for different *P. gingivalis* strains on BSA-MM. **b**, Representative growth curves for mutant *ragAB* variants (n = 3, mean ± standard deviation). **c,d**, Gene expression levels determined by qPCR (n = 3, mean ± standard deviation). **e**, Outer membrane protein expression levels for RagAB variants from W83 wild type by SDS-PAGE densitometry, showing expression levels of RagAB mutants (n = 2, mean ± standard deviation). **f**, SDS-PAGE gel showing OM protein expression levels following removal of inner membrane proteins by sarkosyl treatment. **g**, Representative growth curves (n = 3) for growth on BSA-MM for wild type *P. gingivalis* W83 and ATCC 33277 and strains in which RagAB or RagB from ATCC 33277 was replaced with the corresponding ortholog from W83. **h**, Gene expression levels determined by qPCR. **i**, OM protein expression levels of RagAB.

Compared to wild type W83 RagAB, growth of the *ragA* and *ragAB* deletion strains exhibits a clear phenotype characterised by long (~15-20 hrs) lag periods (Fig. 5b). Unlike the gingipain-*null* KRAB strain^11^, both mutant strains grow eventually, most likely due to passive, RagAB-independent uptake of small peptides produced after prolonged digestion of BSA by gingipains or other *P. gingivalis* proteases. Interestingly, the Δ*ragB* strain grows reasonably well on BSA (Fig. 5b). Since the strain still produces RagA (albeit at low levels; Figs. 5e,f) this demonstrates that RagB, in contrast to RagA, is not required for growth on BSA *in vitro*^12^. Collectively, these data show that RagAB is important for uptake of extracellular peptides produced by gingipains, and provide an explanation for data showing that the transporter is important for the *in vivo* fitness and virulence of *P. gingivalis*^20,21^.

We next constructed several mutant strains for structure-function studies. For RagA, the following mutants were made (see Methods for details): a Ton box deletion (ΔTonB), a DUF domain deletion (ΔDUF), a putatively monomeric RagAB version (RagAB_mono_) which should prevent RagA_2_B_2_ complex formation, and deletions of the two RagA loops (L7 and L8) that remain associated with RagB during lid opening (Δhinge1 and 2). For RagB, a deletion of the acidic loop was made (Δ_AL_), and the acidic loop was converted into a basic loop (RagB_BL_). To complement the growth curves we also determined gene expression levels by qPCR (Figs. 5c,d) and OM protein expression levels via SDS-PAGE (Figs. 5e,f). The ΔTonB and RagAB_mono_ strains have a similar phenotype as ΔRagAB and ΔRagA, suggesting they cannot take up oligopeptides produced by gingipains. However, while the gene expression levels of all variants except the deletions vary only up to ~2-fold (Fig. 5d), the OM protein profiles show that RagAB_mono_ is expressed at very low levels (Figs. 5e,f). On the other hand, OM levels of the ΔTonB mutant are similar to that of wild type, demonstrating that this variant is truly inactive and therefore that RagAB is a *bona-fide* TBDT. For both RagA hinge loop mutants, OM expression levels are very low, so that no conclusions about functionality can be drawn. The RagB acidic loop mutants show intermediate growth on BSA-MM, suggesting they have impaired oligopeptide uptake. However, the OMP profiles of the acidic loop mutants are substantially lower than wild type (Fig. 5e), suggesting that both acidic loop mutants are likely functional, and that the striking inability of *rag-4 P. gingivalis* strains (*e.g*. ATCC 33277) to grow on BSA (Fig. 5a) is not due to the absence of the RagB acidic loop. The most intriguing result was obtained for the RagA ΔDUF mutant, which resembles the KRAB strain (Fig. 5b) and does not grow even after prolonged time, despite being expressed at reasonable levels (Figs. 5e,f). Since the ΔRagA and ΔRagAB strains (which also lack the DUF) do grow after a lag phase, the results suggest an intriguing and important role for the DUF.

To further investigate differences in substrate specificities between the RagAB complexes from W83 and ATCC 33277 we constructed an ATCC 33277 strain in which W83 *ragAB* was expressed from a single-copy plasmid in the ATCC *ΔragAB* background. We also made a strain in which the genomic copy of ATCC *ragB* was replaced by W83 *ragB*. Remarkably, replacement of either RagB or RagAB from ATCC with the corresponding orthologs from W83 results in robust growth of the ATCC strain on BSA-MM (Figs. 5g-i). These results have several important implications. First, they confirm that RagAB is essential for growth on extracellular protein-derived peptides. Second, given the high sequence identities for RagA (~70%) and RagB (~50%) from both strains (Supplementary Fig. 1), it is likely that relatively small differences in transporter structure substantially affect substrate specificity and transport. Third, and perhaps most surprisingly, RagB alone is sufficient to change the substrate specificity of the complex, demonstrating that different RagB lipoproteins can form functional complexes with the same RagA transporter.

### RagAB binds oligopeptides *in vitro* and *in vivo*

Having shown that RagAB is required for growth on peptides derived from extracellular proteins, we next characterised peptide binding to RagAB in more detail. This was complicated by the fact that it was completely unclear what peptide sequences to test. The only clue came from a recent study where an 11-residue peptide of the arginine deiminase ArcA from *Streptococcus cristatus* was proposed to bind to RagB from ATCC 33277^22^. This peptide, designated P4 by the authors, has the sequence NIFKKNVGFKK, and we reasoned that its highly basic character (+4 net charge) combined with the acidic loop in W83 RagB might make it also a good substrate for W83 RagAB. Initial isothermal titration calorimetry (ITC) experiments showed that P4 addition to buffer without protein generated very large heats, precluding ITC as a method to assess peptide binding.

We next carried out microscale thermophoresis (MST) using N-terminally fluorescein-labelled P4 peptide (P4-FAM) and unlabelled RagAB, and obtained dissociation constants of ~2 μM for W83 RagAB and approximately 0.2 μM for RagAB from ATCC 33277 (Figs. 6a,b). The negative control OM protein complex Omp40-41 did not show any binding (Fig. 6c), demonstrating that these results are not due to non-specific partitioning of the labelled peptide into detergent micelles driven by the large fluorescent group.

**Figure 6.**
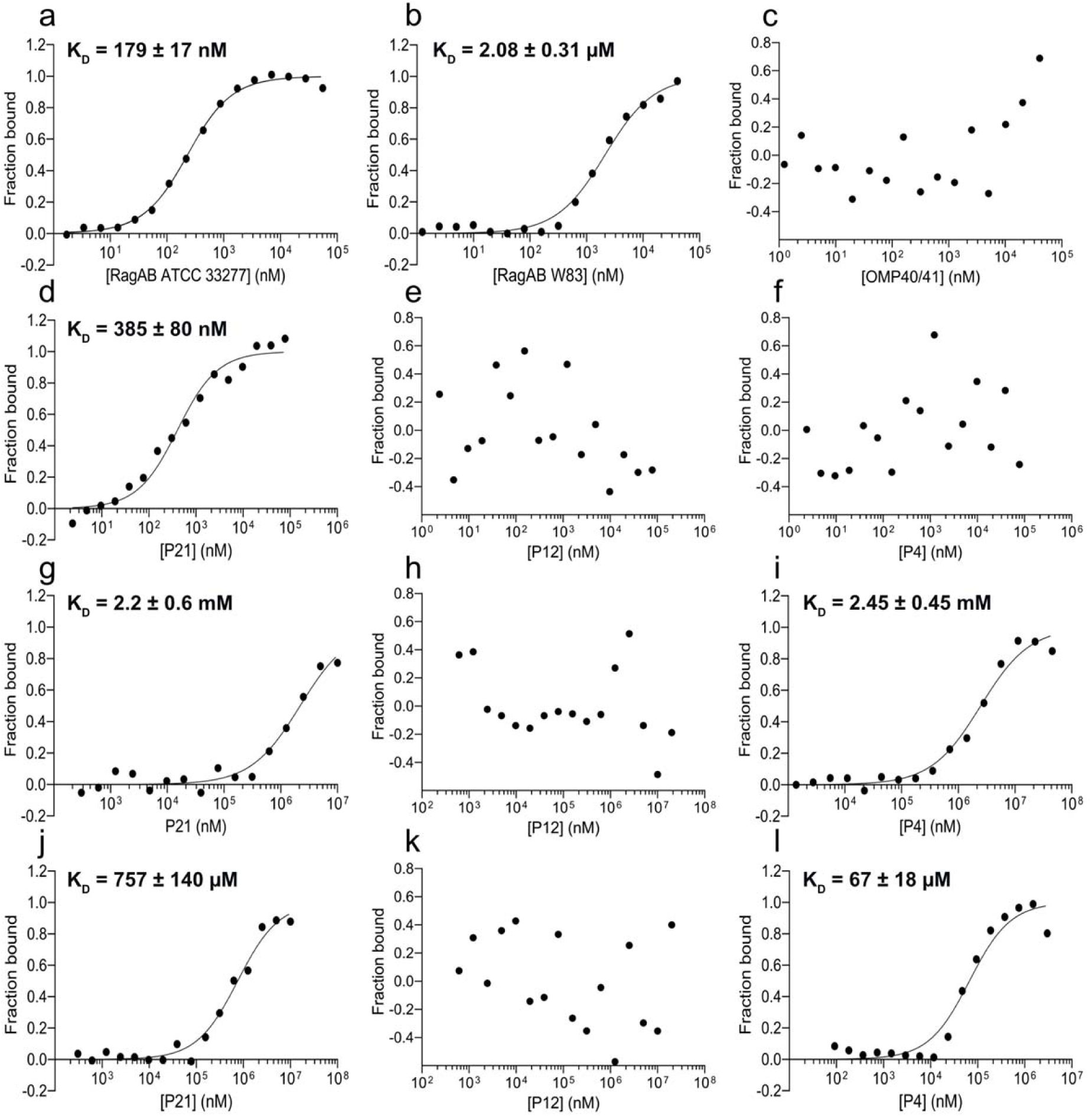
RagAB and RagB bind peptides selectively. MST titration curves with the P4-FAM peptide for (**a**) RagAB from ATCC 33277, (**b**) RagAB from W83 and (**c**) Omp40-41 from W83 (negative control). **d-l**, MST profiles for unlabelled P21, P12 or P4 binding to His-tag labelled W83 RagAB (**d-f**), His-tag labelled W83 RagB (**g-i**), and His-tag labelled ATCC 33277 RagB (**j-l**). Experiments and listed K_d_ values represent the mean of three independent experiments ± SD.

### Identification of RagAB-bound peptides

To identify peptides bound to purified RagAB we employed a sensitive LC-MS/MS peptidomics approach (Methods). As a negative control we again used Omp40-41. Purified RagAB from W83 KRAB was associated with several hundred unique peptides (Supplementary Table 3), whereas none were present in the Omp40-41 sample. Surprisingly, no P4 peptide was detected. All peptides originated from endogenous, mostly abundant *P. gingivalis* proteins representing all cellular compartments, such as ribosomal proteins (L7/L12, L29, S10), OM proteins (RagA, Omp40-41) and inner membrane proteins (signal peptide peptidase). These data suggest that P4 either didn’t bind strongly and/or was outcompeted by excess endogenous *P. gingivalis* peptides after cell lysis. As a comparison, we also analysed peptides bound to RagAB purified from the gingipain-expressing W83 wild-type strain. Perhaps unsurprisingly due to dominant gingipain activity post-cell lysis, many RagAB-associated peptides from wild-type W83 were derived from different proteins compared to the KRAB strain (*e.g*. elongation factor Tu). Almost all peptides from the wild-type dataset have either Lys or Arg at the C-terminus, consistent with the combined trypsin-like specificity of gingipains. Only a minority of peptides from the KRAB strain have a C-terminal basic residue, consistent with the lack of gingipain activity in this strain.

Any in-depth analysis of RagAB-bound peptides is difficult because of the non-quantitative nature of MS when comparing different peptides. The peptides bound to RagAB from W83 KRAB vary in length from 7 to 29 residues, with a broad maximum of around 13 residues (Fig. 7a) that fits well with the peptide density observed in the structures. Assuming equal abundance of each detected peptide, there is a slight preference for neutral to slightly acidic peptides, and the pI distribution has a bimodal shape, with maxima for acidic and slightly basic peptides (Figs. 7b,c). Analysis of the smaller RagAB-bound peptide set from wild-type W83 yields a slightly wider size range from 5-36 residues (Fig. 7d), but overall there are no dramatic differences between the RagAB-bound peptide populations from W83 KRAB and wild-type strains (Figs. 7e,f). Analysis of peptide amino acid frequency shows a substantial enrichment of Ala, Glu, Lys, Thr and Val. By contrast, aromatics (Phe, Trp) and bulky hydrophobics (Leu) appear to be under-represented (Fig. 7g). A semi-quantitative treatment of the data in which the number of times a particular peptide is observed is used as a proxy for relative abundance, yields similar conclusions for the KRAB strain. For the wild-type W83 dataset, a pronounced shift towards basic peptides is observed due to the high abundance of peptides from the N-terminus of elongation factor Tu (Supplementary Fig. 4).

**Figure 7.**
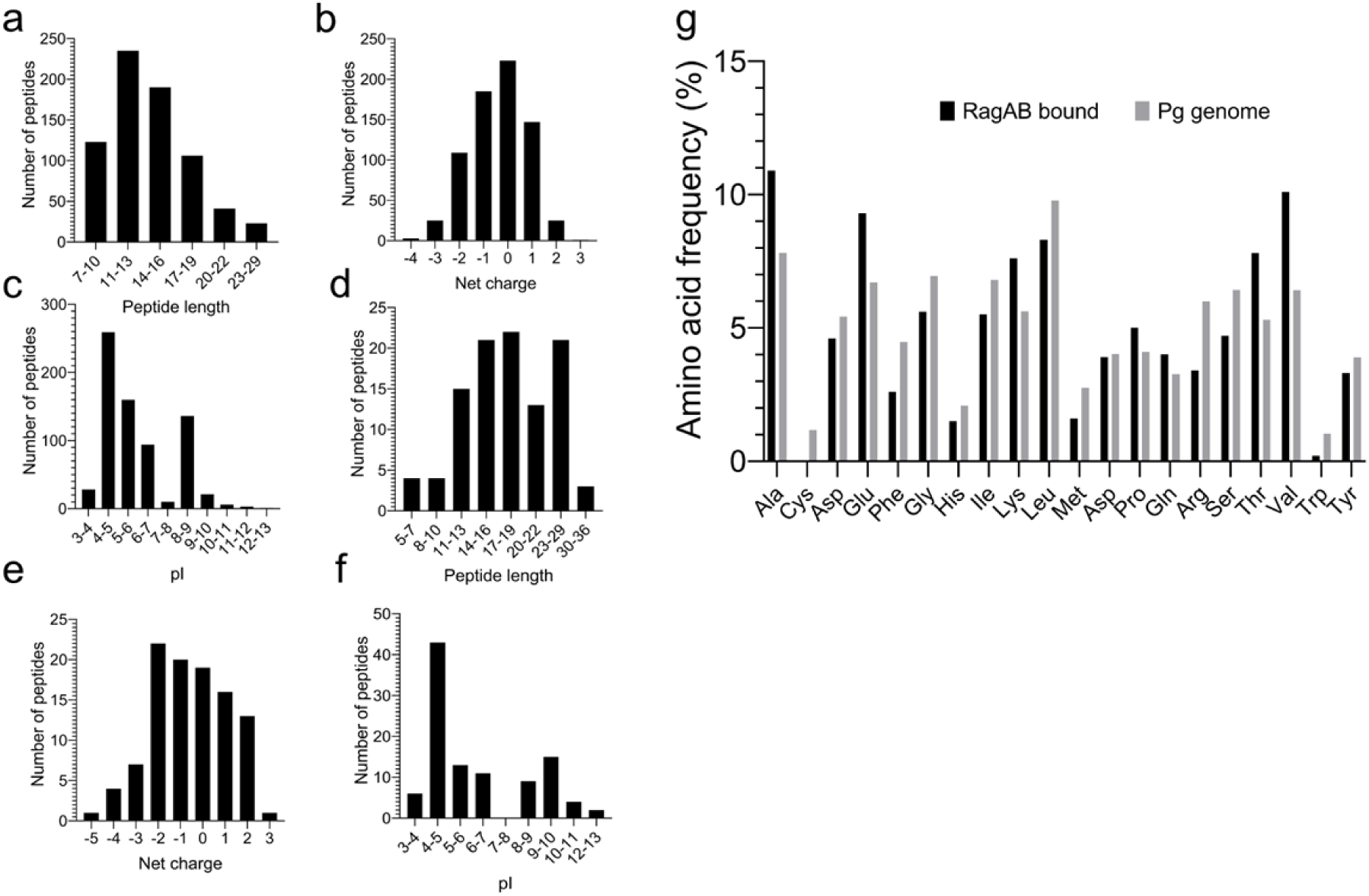
Peptidomics of RagAB. **a-c**, LC-MS/MS analysis of peptides bound to RagAB W83 KRAB, showing length distribution (**a**), total charge (**b**) and pI (**c**). **d-f**, Analysis of peptides bound to RagAB W83 wild-type, showing length distribution (**d**), total charge (**e**) and pI (**f**). **g**, Amino acid frequency of RagAB-bound peptides (KRAB and wild-type combined; black) vs. the amino acid composition in the *P. gingivalis* proteome (gray).

### Peptide binding by RagAB is selective

Based on chromatogram peak heights and the number of MS/MS spectra observed for a particular peptide, we identified the 21-residue peptide KATAEALKKALEEAGAEVELK (henceforth named P21; charge -1) from the C-terminus of ribosomal protein L7 as being very abundant in W83 KRAB RagAB. The P21 peptide was synthesised in addition to its 12-residue core sequence (DKATAEALKKAL, denoted P12; charge +1) that is also present in a number of similar peptides but, crucially, is not identified as a RagAB-bound peptide. We next performed MST experiments on P21, P12 and P4, using unlabelled peptides and His-tag labelled W83 RagAB. Remarkably, we observed robust binding only for P21 (K_d_ ~0.4 μM; Fig. 6d). While the fit to a single binding site model is reasonable, fitting to a model assuming two non-equivalent sites yields lower residuals (Supplementary Fig. 5). This provides the first indication that the two binding sites in RagA_2_B_2_ may not be equivalent, perhaps due to cooperativity between the two RagAB transporter units. The results for P12 and P4 (Figs. 6e,f), suggests that these peptides are unable to displace the bound endogenous peptides. The result for unlabelled P4 contrasts with that for P4-FAM (Fig. 6b), and suggests that the binding of P4-FAM may be driven by the fluorescein moiety. We next asked whether the three peptides bind to RagB. W83 RagB produced good-quality binding curves with P21 and P4, with similar dissociation constants of ~2 mM, but no binding was observed for P12 (Figs. 6g-i). ATCC RagB bound P21 with similar affinity (K_d_ ~0.7 mM) as W83 RagB. In agreement with its recent identification from peptide array analysis^22^, P4 binds well to ATCC RagB, with a dissociation constant of ~70 μM (Figs. 6g-i). Again, no binding was observed for P12 (Fig. 6k). The P21 data suggest that peptides bind with much lower affinities to RagB (P21 K_d_ ~2 mM) compared to RagAB (P21 K_d_ ~0.4 μM), which makes sense assuming that after initial capture by RagB, the peptide needs to be transferred to RagA. The data also show that the P4 and P12 peptides are not good substrates for W83 RagAB and demonstrate that the transporter has a considerable degree of substrate selectivity.

Since we can measure peptide binding to purified RagAB *in vitro* by MST, added peptides compete with co-purified endogenous peptides. Together with robust growth observed in BSA-MM we asked whether we could detect acquisition of BSA tryptic peptides by RagAB *in vitro* (Methods). Indeed, the sample incubated with the BSA digest revealed six bound BSA peptides in addition to endogenous peptides, suggesting that the BSA peptides only partly replace the co-purified ensemble (Supplementary Table 3). This can be explained by the fact that the BSA digest contains about 65 different peptides (Supplementary Table 3), such that the concentration of the BSA “binder” peptides is not high enough to replace all endogenous peptides. By contrast, a 100-fold excess of P21 completely displaced the co-purified peptides (Supplementary Table 3). The fact that only a subset of BSA peptides binds to RagAB reinforces our notion that the transporter is selective. We also incubated the BSA tryptic digest experiment with purified W83 RagB. Prior to incubation, RagB contains only one bound co-purified peptide. After incubation and post-SEC, two BSA peptides are detected, demonstrating that at least some peptides bind to RagB with sufficient affinity to survive SEC (Supplementary Table 3).

To build a unique peptide sequence in the electron density maps we co-crystallised RagAB W83 in the presence of a 50-fold molar excess of P21. Comparison with the original maps obtained from RagAB W83 KRAB shows peptide density at the same site, but with substantial differences, in particular at the N-terminus and positions 9 and 10 of the peptide (Supplementary Fig. 6). However, given that different peptide ensembles co-purify in transporters derived from W83 KRAB and wild-type strains we also crystallised RagAB purified from wild-type W83. The densities for the peptide ensembles from W83 KRAB and wild-type strains are similar, with reasonable fits for the same 13-residue peptide model (ASTTGANSQRGSG). By contrast, the peptide model for the P21 co-crystallisation structure has to be changed substantially for a good fit. The P21 sequence does not fit the density unambiguously, and we speculate that P21 and other peptides are bound with register shifts and perhaps different chain directions. Despite this, in all three structures the same RagAB residues hydrogen bond with the backbones of the modelled substrates, providing a clear rationale how the transporter can bind many oligopeptides.

Our combined data show that RagAB is a dynamic OM oligopeptide transporter that is important for growth of *P. gingivalis* on extracellular protein substrates and possibly for peptide-mediated signalling^22^. We propose a transport model in which the open RagB lid binds substrates followed by delivery to RagA via lid closure (Supplementary Fig. 7a). This would make the RagA Ton box accessible for interaction with TonB, followed by formation of a transport channel into the periplasmic space. To test the premise that substrate binding induces lid closure we collected cryo-EM data on RagAB in the absence and presence of excess P21. Particle classification indeed shows a decrease in OO states and a clear increase in CC states in the P21 sample, in accordance with our model (Supplementary Figs. 7b,c).

## Discussion

How does signalling to TonB occur, and how are unproductive interactions of TonB with “empty” transporters avoided? In the classical, smaller TBDTs such as FecA and BtuB, the ligand binding sites involve residues of the plug domain, leading to the supposition that ligand binding is allosterically communicated to the periplasmic face of the plug. However, there are no direct interactions between the visible, well-defined part of the peptide substrates and the RagA plug. What, then, causes the observed conformational changes in the plug? One possibility is that parts of peptide substrates that are invisible (*e.g*. due to mobility) contact the plug. Given the very large solvent-excluded RagAB cavity even with the modelled 13-residue peptide (~ 9800 Å^3^; ~11500 Å^3^ without peptide; Supplementary Fig. 2), there is enough space to accommodate the long substrates identified by the peptidomics, and these could contact the plug directly. Regardless of these considerations, the presence of substrate density in the open states of the cryo-EM structures suggests that it may not be the occupation of the binding site *per se* that is important for TonB interaction, but closure of the RagB lid (Supplementary Fig. 7). Thus, we hypothesise that certain peptides may bind to RagAB, but do not generate the closed state of the complex and signal occupancy of the binding site to TonB. This may provide an alternative explanation for the different MST results for the P4 and P4-FAM peptide titrations to RagAB (Fig. 6). Given the nature of the MST signal, titrating unlabelled P4 to labelled RagAB is likely to give a signal only if the binding causes a conformational change (*e.g*. lid closure in the case of RagAB). In the “reverse” experiment, the readout is on the labelled peptide (P4-FAM), and the large change in mass upon binding could generate a thermophoretic signal in the absence of any conformational change. Thus, P4 may be an example of a substrate that binds non-productively to RagAB.

A fascinating question is why RagAB (and other SusCD-like transporters) are dimeric. To our knowledge, there are no other TBDTs that function as oligomers, and there is no obvious reason why dimerisation would be beneficial. There are few clues in the structures, but when RagA in, *e.g*., the closed complexes of the CC and OC states is superposed, RagB shows a rigid-body shift of ~3-5 Å, and *vice versa*. A similar trend is observed when the open complexes of the OC and OO states are considered. Moreover, the peptide densities in the OO state are stronger than that of the open complex in the OC state (Fig. 3). Together, this suggests that the individual RagAB complexes could exhibit some kind of cross-talk such as cooperative substrate binding. This notion is supported by the MST data for P21 binding to RagAB, suggesting the presence of two non-identical ligand binding sites and negative cooperativity (Supplementary Fig. 5).

The *rag* locus was probably obtained via horizontal gene transfer of the *sus* operon, as it is located on a pathogenicity island of *P. gingivalis*^21,23,24^. In contrast to environmental and commensal Bacteroidetes that have many SusCD systems dedicated to glycan transport^14^, all *P. gingivalis* strains sequenced to date (57 in total) have only one *ragAB* operon. A single *ragAB* operon is also present in all strains of *Porphyromonas gulae* (12 in total), the pathogenic equivalent of *P. gingivalis* in dogs, and in some other closely related oral species of *Porphyromonas*. The distinct chemical nature of various glycans likely requires transporter diversification, and we speculate that this has underpinned the evolution of multiple SusCD systems for glycan transport in *e.g*. gut *Bacteroides spp*. By contrast, the peptide backbone offers a way to achieve polyspecificity, obviating the need to evolve multiple highly selective peptide transporters. The natural habitat for P. *gingivalis* is dental plaque, a biofilm on the tooth surface immersed in gingival crevicular fluid (GCF), an inflammatory exudate which is dominated by a limited number of abundant plasma proteins such as albumin^25^, and we speculate that this may have contributed to RagAB substrate selectivity. By developing a surface-exposed proteolytic machinery (gingipains) together with a transport system (RagAB) that efficiently imports relatively large peptides, *P. gingivalis* likely has a competitive advantage *in vivo*.

A contribution of RagAB to *P. gingivalis* virulence was first proposed based on the observation that both RagA and RagB are recognized by the sera of periodontitis patients, but not by that of healthy controls^26^. The *rag* locus also contributes to host cell invasion by *P. gingivalis*^27^ and soft tissue damage in a murine abscess model^28^. A definitive assignment of the role of the *rag* locus in virulence is complicated by the fact that clinical strains of *P. gingivalis* carry allelic forms of established virulence factors such as fimbria, that are known to be variably associated with the severity of periodontitis^29^. RagAB occurs in 4 well-defined allelic forms (*rag-1*, *-2*, *-3* and *-4*)^20^. Since *rag-1* locus strains are associated with more severe periodontitis^20^ and are more virulent in mice models of infection^28^ argues that the ability of *rag-1* strains to grow on BSA (which is 70 and 76% identical to mice and human albumin, respectively) adds to the pathogenicity of *P. gingivalis in vivo*. Although the contribution of the different *rag* loci to *P. gingivalis* virulence needs further study it is clear that expression of RagAB is absolutely essential to *P. gingivalis* ATCC33277 (*rag-4* genotype) fitness during epithelial colonization and survival in a murine abscess model^30^. Finally, RagB directly stimulates a major pro-inflammatory response in primary human monocytes^31^ and, therefore, may have a direct role the pathobiology of periodontitis. RagB is also under investigation as a potential vaccine against periodontitis^32,33^.

Hoe does RagAB compare to other peptide transporters? There are no other structures of OM peptide transporters, and RagAB is very different, both structurally and mechanistically, from MFS-type peptide transporters^34^. Comparable to *P. gingivalis* in terms of peptide utilisation are Gram-positive lactic acid bacteria such as *Lactococcus lactis*, which acquire peptides derived from milk proteins such as caseins. In the case of *L. lactis*, peptides are generated by a single surface-exposed protease^35^ and taken up by the oligopeptide permease Opp after delivery by the associated surface-exposed OppA receptor lipoprotein^36^. *L. lactis* OppA binds peptides from 4-35 residues in length, with an optimum for nonapeptides, and is functionally equivalent to RagB. No structure of an Opp transporter is available, and it is not clear how OppA interacts with the transporter and how the peptides are transferred from OppA to the permease. Another difference between the Opp system and RagAB is that OppA consists of two domains that change conformation upon peptide binding, enclosing a very large, solvent-excluded binding cavity^37^. While specific interactions with peptides involve only six residues, OppA binds substrates with high affinity (bradykinin Kd = 0.1 μM^36^), and most likely requires external energy input for peptide release. By contrast, RagB is a single-domain protein that does not undergo major conformational changes upon substrate binding. Together with the relatively flat shape of the substrate binding surface (Fig. 1g) this explains the relatively low affinity of RagB for peptide substrates, enabling their transfer to RagA.

## Author contributions

MM, JP and BvdB initiated the project. MM cultured cells, purified and crystallised proteins and performed MST binding experiments, with guidance from JP and BvdB. JBRW and SR determined cryo-electron microscopy structures, supervised by NR. ZN performed cloning and strain construction. GB carried out qPCR experiments, and CS and JJE performed the peptidomics analyses. KP performed the MD simulations, supervised by UK. BvdB purified and crystallised proteins and determined the RagAB crystal structures. The manuscript was written by BvdB with input from MM, JBRW, NR and JP.

## Acknowledgements

This work was supported by a Welcome Trust Investigator award (214222/Z/18/Z to BvdB). We would further like to thank personnel of the Diamond Light Source for beam time (Block Allocation Group numbers mx-13587 and mx-18598) and assistance with data collection. All EM was performed at the Astbury Biostructure Laboratory which was funded by the University of Leeds and the Wellcome Trust (108466/Z/15/Z). We thank Drs. Rebecca Thompson, Emma Hesketh and Dan Maskell for EM support. We would also like to thank T. Kantyka for help in designing MS experiments. This study was supported in part by National Science Centre, Poland grants UMO-2015/19/N/NZ1/00322 and UMO-2018/28/T/NZ1/00348 to MM, UMO-2016/23/N/NZ1/01513 to ZN, UMO-2018/29/N/NZ1/00992 to GPB, and UMO-2018/31/B/NZ1/03968 and NIDCR/DE 022597 (NIH) to JP. LC-MS/MS analysis was supported by the Novo Nordisk Foundation (BioMS). JBRW was supported by a Wellcome Trust 4 year PhD studentship (215064/Z/18/Z).

## Methods

### Bacterial strains and general growth conditions

*Porphyromonas gingivalis* strains (listed in Table S4) were grown in enriched tryptic soy broth (eTSB per liter: 30 g tryptic soy broth, 5 g yeast extrac; further supplemented with 5 mg hemin; 0.25 g L-cysteine and 0.5 mg menadione) or on eTSB blood agar (eTSB medium containing 1.5% [w:v] agar, further supplemented with 5% defibrinated sheep blood) at 37 °C in an anaerobic chamber (Don Whitley Scientific, UK) with an atmosphere of 90% nitrogen, 5% carbon dioxide and 5% hydrogen. *Escherichia coli* strains (listed in Supplementary Table 4), used for all plasmid manipulations, were grown in Luria–Bertani (LB) medium and on 1.5% agar LB plates. For antibiotic selection in *E. coli*, ampicillin was used at 100 μg/ml. *P. gingivalis* mutants were grown in the presence of erythromycin at 5 μg/ml and/or tetracycline at 1 μg/ml.

### Growth of *P. gingivalis* in minimal medium supplemented with BSA

*P. gingivalis* strains were grown overnight in eTSB. The cultures were washed twice with enriched Dulbecco’s Modified Eagle’s Medium (eDMEM per liter: 10 g BSA; further supplemented with 5 mg hemin; 0.25 g L-cysteine and 0.5 mg menadione) and finally resuspended in eDMEM. The OD_600_ in each case was adjusted to 0.2 and bacteria were grown for 40 hours. OD_600_ was measured at 5 hour intervals.

### Mutant construction

For RagA, the following mutants were made: a Ton box deletion (ΔTonB; residues V100-Y109 deleted), a DUF domain deletion (ΔDUF; residues V25-K99 deleted) a putatively monomeric RagAB version made via introduction of a His6-tag between residues Q570-G571; RagAB_mono_), and deletions of the two loops (L7 and L8) that remain associated with RagB during lid opening as based on the cryo-EM structures (Δhinge1, residues Q670-G691 deleted and replaced by one Gly; Δhinge2, residues L731-N748 deleted and replaced by one Gly). For RagB, two variants were made, both involving the acidic loop: deletion of this loop (Δ_AL_; residues R97-S104) and conversion of the acidic loop into a basic loop (RagB_BL_; D99/E100/D101/E102 replaced by R99/K100/R101/K102). All *P. gingivalis* mutants were prepared using wild-type W83 strain or its derivatives (unless stated otherwise) and constructed by homologous recombination^38^. The “swap” mutants were prepared using the ATCC33277 strain as background. For deletion strains, 1 kb regions upstream (5’) and downstream (3’) from the *ragA*, *ragB* and both *ragAB* genes, as well as chosen antibiotic resistance cassettes, were amplified by PCR. Obtained DNA fragments were cloned into pUC19 vector using restriction digestion method and/or Gibson method^39^. For the construction of master plasmids for RagA and RagB mutants the whole gene sequences were amplified with addition of antibiotic cassettes, 1kb downstream fragments and cloned into pUC19 vector. Desired mutations were introduced into master plasmids by the SLIM PCR method^40^. A similar method was used for the “swap” RagB-W83inATCC and RagB-ATCCinW83 strains. In both deletion RagB plasmids we inserted the gene of opposite strain (i.e. *ragB* from W83 into ΔRagB-ATCC). We also obtained a RagAB “swap” strains using pTIO-1 plasmid^41^. The DNA sequences of *ragAB* from both strains were cloned with the addition of their promoters into pTIO plasmids and further conjugated with the opposite deletion strains using *E. coli* S-17 λpir (i.e. RagAB-W83-pTIO into delRagAB-ATCC)^42^. Primers used for plasmid construction and mutagenesis are listed in Supplementary Table 4. All plasmids were analyzed by PCR and DNA sequencing. *P. gingivalis* competent cells^43^ were electroporated with chosen plasmids and plated on TSBY with appropriate antibiotics – erythromycin (5 µg/ml) or tetracycline (1 µg/ml) and grown anaerobically for approximately 10 days. Clones were selected and checked for correct mutations by PCR and DNA sequencing. Bacterial strains generated and used in this study are listed in Supplementary Table 4.

### W83 KRAB RagAB production and purification

The (non-His tagged) RagAB complex from *P. gingivalis* W83 KRAB was isolated from cells grown in rich media. In brief, cells from 6 l of culture were lysed by 1 pass through a cell disrupter (0.75 kW; Constant Biosystems) at 23,000 psi, followed by ultracentrifugation at 200,00 × g for 45 minutes to sediment the total membrane fraction. The membranes were homogenised and pre-extracted with 100 ml 0.5% sarkosyl in 20 mM Hepes pH 7.5 (20 min gentle stirring at room temperature^44^) followed by ultracentrifugation (200,000 × g; 30 min) to remove inner membrane proteins. The sarkosyl wash step was repeated once, after which the pellet (enriched in OM proteins) was extracted with 100 ml 1% LDAO (in 10 mM Hepes/50 mM NaCl pH 7.5) for 1 hour by stirring at 4 °C. The extract was centrifuged for 30 min at 200,000 × g to remove insoluble debris. The solubilised OM was loaded on a 6 ml Resource-Q column and eluted with a linear NaCl gradient to 0.5 M over 20 column volumes. Fractions containing RagAB were ran on analytical SEC (Superdex 200 Increase GL 10/300) in 10 mM Hepes/100 mM NaCl/0.05% LDAO pH 7.5 in order to obtain RagAB of sufficient purity. Finally, the protein was detergent-exchanged to C_8_E_4_ using two rounds of ultrafiltration (100 kDa MWCO), concentrated to 15-20 mg/ml and flash-frozen in liquid nitrogen.

### Wild type W83 RagAB production and purification

Homologous recombination was used to add a 8×His-tag to the C terminus of genomic *ragB* in the *P. gingivalis* W83 strain. The mutant was grown about 20 h in rich medium under anaerobic conditions. The cells from 6 l of culture were collected and processed as outlined above. The insoluble material was homogenised with 1.5% LDAO (in 20 mM Tris-HCl/300 mM NaCl pH 8.0) and the complex was purified by nickel-affinity chromatography (Chelating Sepharose; GE Healthcare) followed by gel filtration using a HiLoad 16/60 Superdex 200 column in 10 mM Hepes/100 mM NaCl, 0.03% DDM pH 8.0.

### Purification of RagB from W83 and ATCC 33277 expressed in *E. coli*

Genes encoding for the mature parts of RagB from W83 and ATCC 33277 (with His6-tags at the C-terminus) were amplified by PCR from genomic DNA extracted from *P. gingivalis*. The DNA fragments were purified and cloned into the arabinose-inducible pB22 expression vector^45^ using NcoI/XbaI restriction enzymes. The obtained expression plasmid was transformed into *E. coli* strain BL21 (DE3). Transformed *E. coli* cells were grown in LB media containing ampicillin (100 μg/ml) at 37 °C to an OD_600_ ~ 0.6 and expression of the recombinant protein was induced with 0.1% arabinose. After ~2.5 h at 37 °C, cells were collected by centrifugation (5,000 × g; 15 min), resuspended in 20 mM Tris-HCl/300 mM NaCl pH 8.0 and lysed by 1 pass through a cell disrupter (0.75 kW; Constant Biosystems) at 23,000 psi, followed by ultracentrifugation at 200,000 × g for 45 minutes. The supernatant was loaded on nickel-affinity resin (Chelating Sepharose; GE Healthcare) and after washing with 30 mM imidazole, protein was eluted with buffer containing 250 mM imidazole. Protein was further purified by gel filtration in 10 mM Hepes/100 mM NaCl pH 7.5 using a HiLoad 16/60 Superdex 200 column.

### Crystallisation and structure determination of RagAB from W83 KRAB

Sitting drop vapour diffusion crystallisation trials were set up using a Mosquito crystallisation robot (TTP Labtech) using commercially available crystallisation screens (MemGold 1 and 2; Molecular Dimensions). Initial hits were optimised manually by hanging drop vapour diffusion. Crystals were cryoprotected by transferring them for 5-10 s in mother liquor containing an additional 10% PEG400. A few crystals optimised from MemGold 2 condition C8 (18% PEG200, 0.1 M KCl, 0.1 M K-phosphate pH 7.5) diffracted anisotropically to below 4 Å at the Diamond Light Source (DLS) synchrotron at Didcot, UK (space group C222_1_; cell dimensions ~190 × 377 × 369 Å, with four RagAB complexes in the asymmetric unit). Data were processed via Xia2^46^ or Dials^47^. The structure was solved by molecular replacement with Phaser^48^, using data to 3.4 Å resolution. The structures of RagB (PDB 5CX8) and a Sculptor-modified model of BT2264 SusC (PDB 5FQ8) were used as search models. The RagAB model was built iteratively by a combination of manually building in COOT^49^ and the AUTOBUILD routine within Phenix^50^, and was refined with Phenix^51^ using TLS refinement with 1 group per chain. The final R and R_free_ factors of the RagAB structure are 20.6 and 25.3%, respectively (Supplementary Table 1). Structures of RagAB purified from wild type W83 (+/- P21) were solved via molecular replacement using Phaser, using the best-defined RagAB complex as search model. Structures were refined within Phenix as above (Supplementary Table 1) and structure validation was carried out with MolProbity^52^.

### Crystallisation and structure determination of wild type RagAB W83 in the absence and presence of P21

Crystallisation trials were performed as outlined above with the following modifications: the protein was not detergent-exchanged after gel filtration; two other commercially available crystallisation screens were used (MemChannel and MemTrans; Molecular Dimensions); For co-crystallisation of RagAB W83 with P21 peptide, RagAB W83 at a concentration of 17 mg/ml (~0.1 mM) was incubated overnight at 4 °C with 5 mM P21 peptide. For RagAB W83 the crystals were optimised from MemTrans condition F6 (22% PEG400, 0.07 M NaCl, 0.05 M Na-citrate pH 4.5), for RagAB W83 + P21 the crystals were optimised from MemChannel condition D3 (15% PEG1000, 0.05 M Li-sulfate, 0.05 M Na-phosphate monobasic, 0.08 M citrate pH 4.5). Crystals were cryoprotected by transferring them for 5-10 s in mother liquor containing additional PEG400 to generate a final concentration of ~25%.

### Microscale thermophoresis

The Monolith NT.115 instrument (NanoTemper Technologies GmbH, Munich, Germany) was used to analyse the binding interactions between P4, P12 and P21 peptides and RagAB from W83 as well as RagB from W83 and ATCC 33277. Proteins were labelled with Monolith His-tag Labeling Kit RED-tris-NTA 2nd Generation (NanoTemper Technologies). For all interactions the concentrations of both fluorescently labelled molecule and ligand were empirically adjusted using Binding Check mode (MO.Control software, NanoTemper Technologies). Experiment was performed in assay buffer (10 mM Hepes/100 mM NaCl pH 7.5) with addition of 0.03% DDM in the case of RagABs. For the measurement, sixteen 1:1 ligand dilutions were prepared and then were mixed with one volume of labelled protein followed by loading into Monolith NT.115 Capillaries. Initial fluorescence measurements followed by thermophoresis measurement were carried out using 100% LED power and medium MST power, respectively. Data for three independently pipetted measurements were analysed (MO.Affinity Analysis software, NanoTemper Technologies) allowing for determination of dissociation constant (K_D_). The data was presented using GraphPad Prism 8. The interaction between unlabelled W83 and ATCC 33277 RagAB and FAM-labelled P4 peptide was done in the same way.

### Isolation of the outer membrane fractions

Outer membrane fractions from 1 l of culture were isolated using sarkosyl extraction method (see RagAB W83 KRAB production and purification). The samples in 10 mM Hepes/50 mM NaCl pH 7.5 containing 1% LDAO were diluted 3 times and loaded on SDS-PAGE. The bands were analysed quantitatively using Image Lab 6.0.1 software (BIO-RAD). The band at ~70 kDa was used as a reference sample (loading control).

### Generation of BSA tryptic mixture

BSA was solubilized in 100 mM ammonium bicarbonate/8M urea pH 8.0. DTT was added to a final concentration of 10 mM and the mixture was incubated for 60 min at RT. Protein was alkylated by addition of iodoacetamide to a final concentration of 30 mM and incubation for 60 min at RT in the dark. Next, the concentration of DTT was adjusted to 35 mM followed by dilution of the sample to 1 M urea. For cleavage, trypsin from bovine pancreas (Sigma) was added at 1:25 (trypsin : BSA) mass ratio and the sample was incubated overnight at 37 °C. Digestion was stopped by acidifying the sample to pH < 2.5 with formic acid. Peptides were purified using Peptide Desalting Spin Columns (Thermo Scientific) and dried using Speed Vacuum Concentrator Savant SC210A (Thermo Scientific).

### Acquisition of BSA-derived peptides and P21 peptide by RagAB and RagB *in vitro*

W83 RagAB and W83 RagB were incubated overnight at 4 °C with an ~100-fold molar excess of tryptic BSA digest (see Generation of BSA tryptic mixture) and P21 peptide. The samples were then ran on Superdex 200 Increase GL 10/300 in 10 mM Hepes/100 mM NaCl pH 7.5 with addition of 0.03% DDM for RagAB W83. Fractions containing protein were pooled, acidified by addition of formic acid to a final concentration 0.1% and analysed by MS.

### qPCR

Samples of 1 ml of bacterial cultures (OD_600_ = 1.0) were centrifuged (5,000 × g; 5 min) at 4°C, pellets were resuspended in 1 ml Tri Reagent (Ambion), incubated at 60 °C for 20 minutes, cooled to room temperature and total RNA was isolated according to manufacturer’s instructions. Genomic DNA was removed from samples by digestion with DNAse I (Ambion); 2 µg of RNA was incubated with 2 U of DNAse for 60 minutes at 37 °C. Following digestion, RNA was purified using Tri Reagent. Reverse transcription of 50 ng of RNA was performed with High-Capacity cDNA Reverse Transcription Kit (Life Technologies), and reaction mixture was then diluted 20 times. Real-time PCR was done in 10 µl reaction volume, using KAPA SYBR FAST qPCR Master Mix (Kapa Biosystems) with 2 µl of diluted reverse transcription mixture as template. Primers used are listed in Table S4. Reaction conditions were 3 minutes at 95 °C, followed by 40 cycles of denaturation for 3 seconds at 95 °C and annealing/extension for 20 seconds at 60 °C. The reaction was carried out with a CFX96 thermal cycler (Bio-Rad), and data was analysed in Bio-Rad CFX Manager software.

### CryoEM sample preparation and data collection

A sample of purified RagAB solubilised in a DDM-containing buffer (10 mM HEPES pH 7.5, 100 mM NaCl, 0.03 % DDM) was prepared at 1.75 mg/ml (principle dataset) or 3 mg/ml (P21 addition experiment). For the P21 addition experiment, control and P21-doped grids (with 50-fold molar excess of P21 peptide) were prepared at the same time from the same purified stock of RagAB for consistency. In all cases, a 3.5 μL aliquot was applied to holey carbon grids (Quantifoil 300 mesh, R1.2/1.3), which had been glow discharged at 10 mA for 30 s before sample application. Blotting and plunge freezing were carried out using a Vitrobot Mark IV (FEI) with chamber temperature set to 6 °C and 100 % relative humidity. A blot force of 6 and a blot time of 6 s were used prior to vitrification in liquid nitrogen-cooled liquid ethane.

Micrograph movies were collected on a Titan Krios microscope (Thermo Fisher) operating at 300 kV with a GIF energy filter (Gatan) and K2 summit direct electron detector (Gatan) operating in counting mode. Data acquisition parameters for each data set can be found in Supplementary Table 2.

### Image processing

Image processing was carried out using RELION (v2.1 and v3.0)^53,54^. Drift correction was performed using MotionCor2^55^ and contrast transfer functions were estimated using gCTF^56^. Micrographs with estimated resolutions poorer than 5 Å and defocus values >4 μm were discarded using a python script^57^. For the principle RagAB dataset, particles were autopicked using a gaussian blob with a peak value of 0.3. Control and experimental datasets for the P21 addition experiment were autopicked using the ‘general model’ in crYOLO^58^. In both cases, particles were extracted in 216 × 216 pixel boxes and subjected to several rounds of 2D classification in RELION^53^. 3D starting models were generated *de novo* from the EM data by stochastic gradient descent in RELION. Processing of RagAB control and P21-doped datasets was only taken as far as 3D classification.

For the principle RagAB dataset, three conformational states representing the CC, OC and OO states were apparent in the first round of 3D classification and the corresponding particle stacks were treated independently in further processing. C2 symmetry was applied to both the CC and OO reconstructions. Post-processing was performed using soft masks and yielded reconstructions for the CC, OC and OO states with resolutions of 3.7 Å, 3.7 Å and 3.9 Å respectively, as estimated by gold standard Fourier shell correlations using the 0.143 criterion. The original micrograph movies were later motion corrected in RELION 3.0^54^. Particles contributing to the final reconstructions were re-extracted from the resulting micrographs. Following reconstruction, iterative rounds of per-particle CTF refinement, with beam tilt estimation, and Bayesian particle polishing were employed which improved the resolution of post-processed CC, OC and OO maps to 3.3 Å, 3.3 Å and 3.4 Å respectively.

### Model building into cryoEM maps

Comparing the maps to the crystal structure of RagAB revealed that their handedness was incorrect. Maps were therefore Z-flipped in UCSF Chimera^59^. The crystal structure was rigid-body fit to the CC density map and subjected to several iterations of manual refinement in COOT and ‘real space refinement’ in Phenix^51^. The asymmetric unit was symmetrised in Chimera after each iteration. Starting models for the OC and OO states of the complex were obtained from the CC structure by rigid-body fitting of one or both RagB subunits to their cognate open density in the OC and OO maps respectively. These too were subjected to several iterations of manual refinement in COOT and ‘real space refinement’ in Phenix. Molprobity was used for model validation^52^.

### Molecular Dynamics simulations

The bound peptide in the refined crystal structure of a RagAB “monomer” was removed *in silico* and the system was inserted into a palmitoyloleoyl-phosphatidylethanolamine (POPE) bilayer using the CHARMM-GUI Membrane Builder^60^. The systems were solvated using a TIP3P water box and neutralized by adding the required counter-ions. Simulations were performed using GROMACS 5.1.2^61^ and the all-atom CHARMM36 force fields^62,63^. For the long-range Coulomb interactions, the partice-mesh Ewald (PME) summation method^64^ has been employed with a short-range cutoff of 12 Å and a Fourier grid spacing of 0.12 nm. In addition, the Lennard-Jones interactions were considered up to a distance of 10 Å and a switch function was used to turn off interactions smoothly at 12 Å. Achieved by semi-isotropic coupling to a Parrinello-Rahman barostat^65^ at 1 bar with a coupling constant of 5 ps, the final unbiased simulations were performed in the isothermal-isobaric (NPT) ensemble. A Nosé-Hoover thermostat^66,67^ was used to keep the temperature at 300 K with a coupling constant of 1 ps. A total of three simulations of 2500 ns were carried out for the apo complex with a time step of 2 fs by applying constraints on hydrogen atom bonds using the LINCS algorithm^68^. Similarly, the dimeric complex of RagA_2_B_2_ in the OO state was simulated for 500 ns. Cavity calculations were performed using CASTp^69^.

### Peptide identification by LC-MS

Bound peptides were isolated from purified RagAB by precipitation via addition of trichloroacetic acid to a final concentration of 30% and incubation at 4°C for 2 hrs. Subsequently the peptide containing supernatants were collected by centrifugation at 17.000xg. The isolated peptides were micropurified using Empore™ SPE Disks of C18 octadecyl packed in 10 µl pipette tips.

LC-MS/MS was performed using an EASY-nLC 1000 system (Thermo Scientific) connected to a QExactive+ Hybrid Quadrupole-Orbitrap Mass Spectrometer (Thermo Scientific). Peptides were dissolved in 0.1 % formic acid and trapped on a 2 cm ReproSil-Pur C18-AQ column (100 μm inner diameter, 3 µm resin; Dr. Maisch GmbH, Ammerbuch-Entringen, Germany). The peptides were separated on a 15-cm analytical column (75 μm inner diameter) packed in-house in a pulled emitter with ReproSil-Pur C18-AQ 3 μm resin (Dr. Maisch GmbH, Ammerbuch-Entringen, Germany). Peptides were eluted using a flow rate of 250 nl/min and a 20-minute gradient from 5% to 35% phase B (0.1% formic acid and 90% acetonitrile or 0.1 % formic acid, 90 % acetonitrile and 5% DMSO). The collected MS files were converted to Mascot generic format (MGF) using Proteome Discoverer (Thermo Scientific).

The data were searched against the *P. gingivalis* proteome (UniRef at uniprot.org) or the Swiss-prot database using a bovine taxonomy. Database search were conducted on a local mascot search engine. The following settings were used: MS error tolerance of 10 ppm, MS/MS error tolerance of 0.1 Da, and either non-specific enzyme or trypsin.

