## Supplementary Information for "Dynamic oligopeptide acquisition by the RagAB transporter from *Porphyromonas gingivalis*"

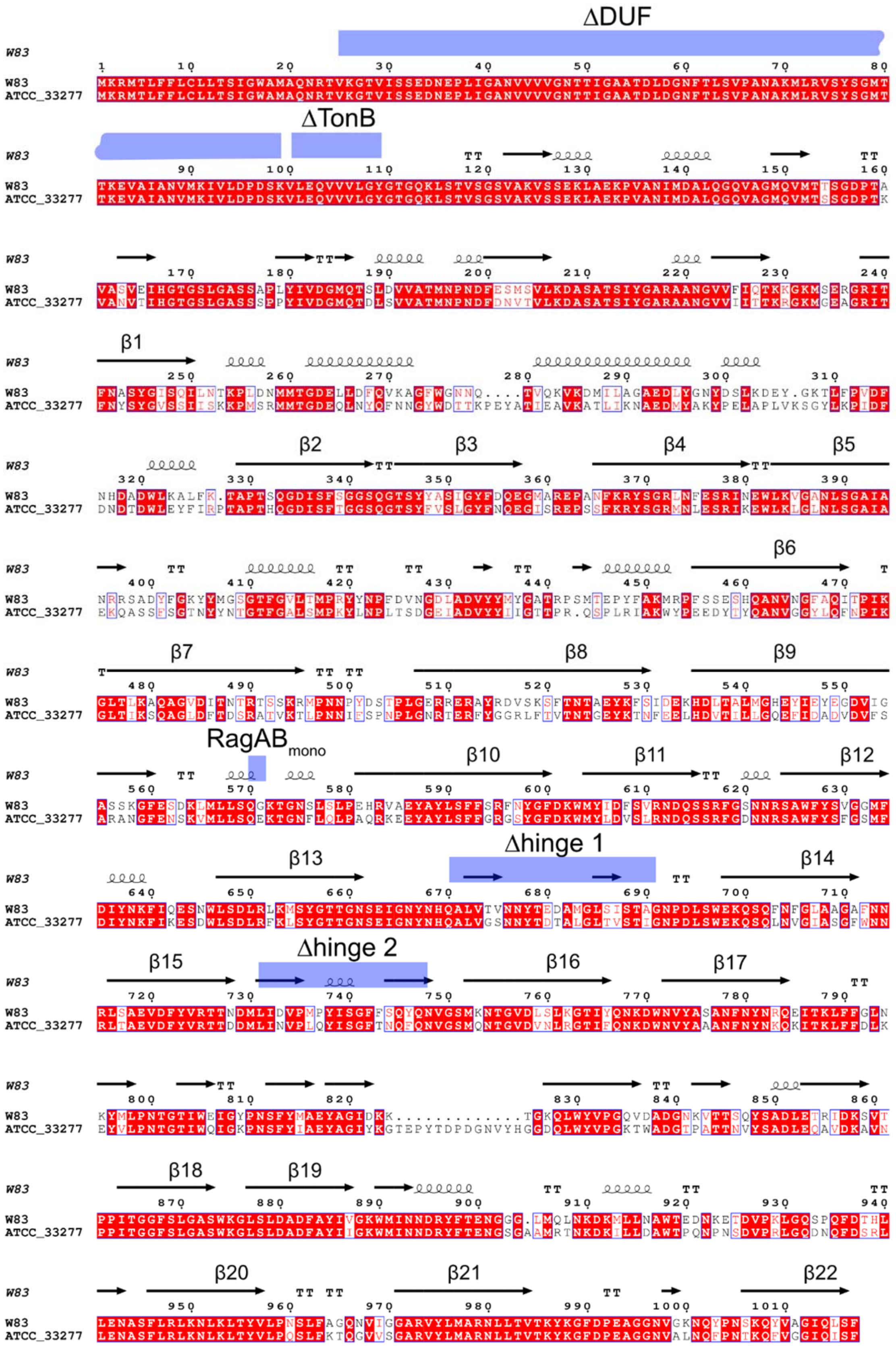

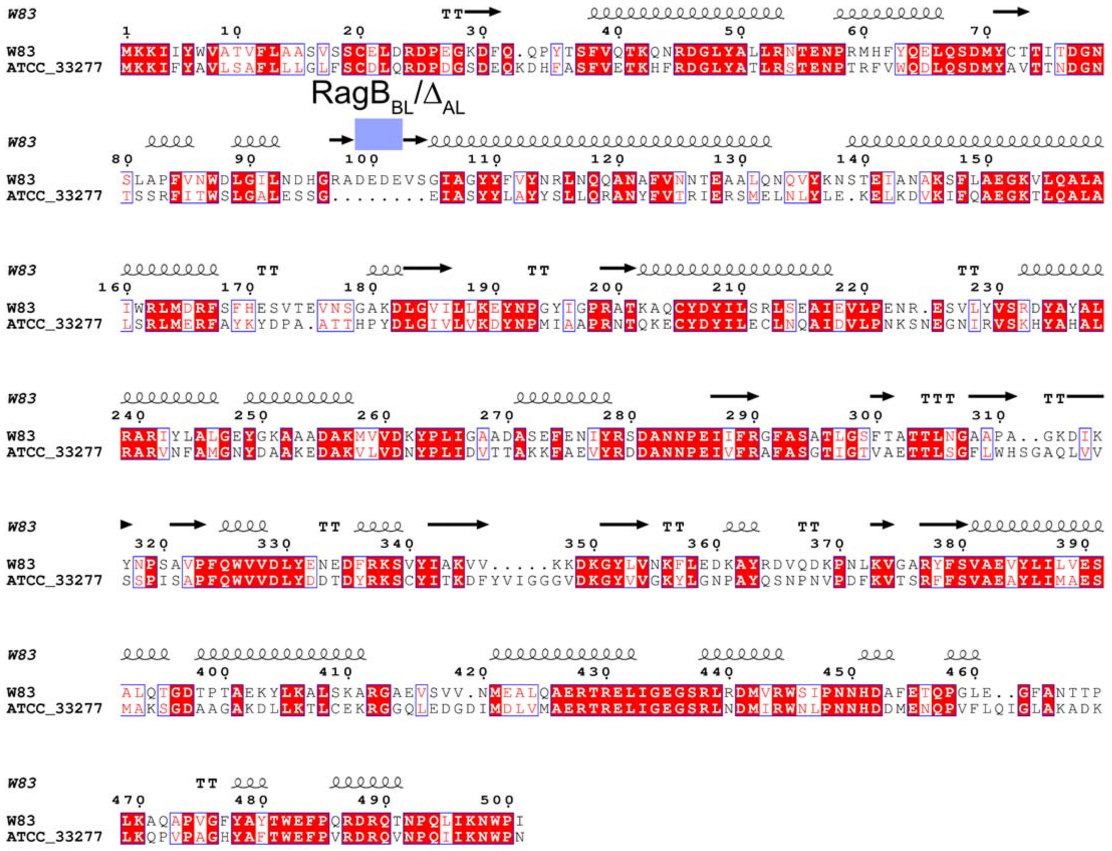


**Supplementary Figure 1 Sequence alignments for RagA (top panel) and RagB (bottom panel) from W83 and ATCC 33277**. The secondary structure assignment based on the RagAB W83 crystal structure is indicated. The positions of the site-directed mutants made are indicated with blue bars. Only the transmembrane β-strands are numbered in RagA. The image was produced by ESPript 3.0[1]. Sequence alignment was performed using Clustal Omega[2].

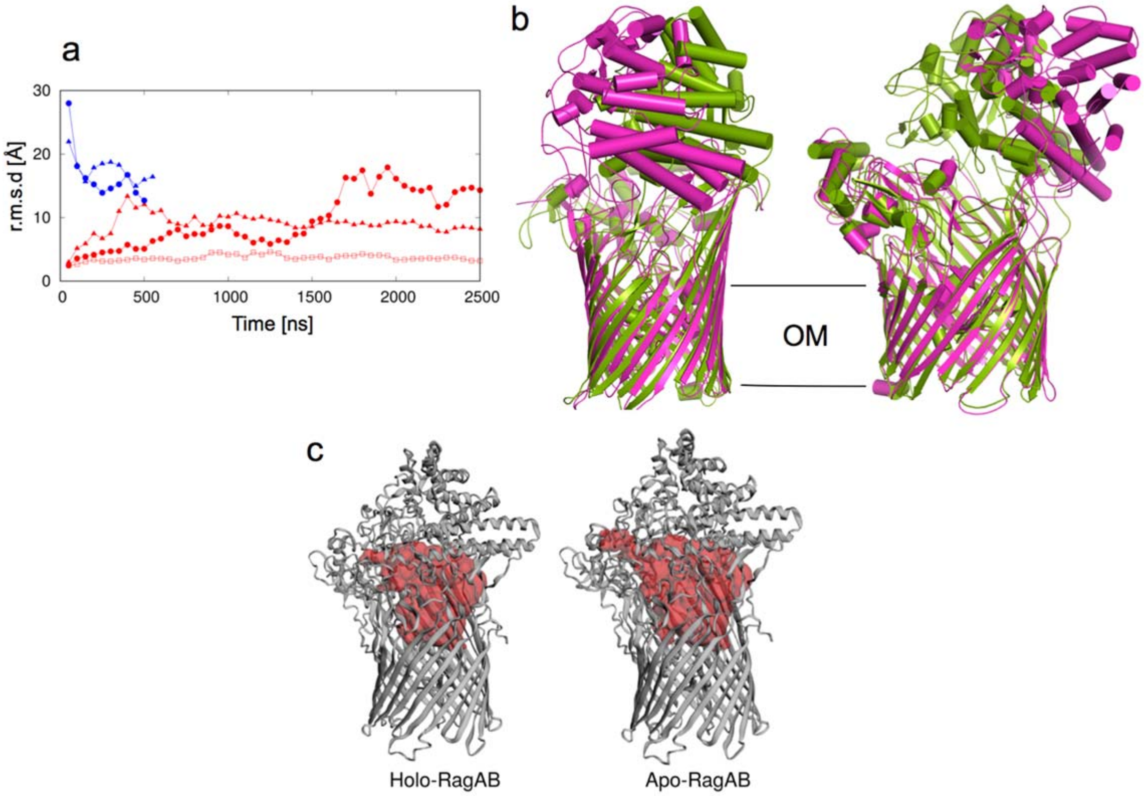


**Supplementary Figure 2 Molecular dynamics simulations of RagAB show lid opening.** **a**, $C_\alpha$-rmsd values of RagB lids in apo-RagAB (red) with reference to the starting crystal structure in the closed conformation. The $C_\alpha$-rmsd values of the RagB lids in $RagA_2B_2$ are shown in blue with reference to the OO EM state. Each point indicates an average of 50 ns simulation trajectory. **b**, Comparison of the $RagA_2B_2$ open conformation from EM (magenta) with the snapshot of the most open simulation at 2500 ns (green). **c**, Internal surface of peptide binding cavities in closed holo-RagAB and apo-RagAB, generated with CASTp[3]. The bound peptide from a RagAB subunit in the crystal structure was removed *in silico* to generate a closed apo-complex, and performed three independent MD simulations. For one of the simulations, a clear opening of the RagB lid was observed, reminiscent of recent results for a SusCD transporter and supporting the notion that ligand removal resets the transporter to favour the open state[4]. The subsequent determination of the EM structures allowed us to compare both open states, and showed that the RagB lid in the simulation opens less wide than that in the EM structure, at least during the timescale of the simulation. We also observed a partial closing of both RagB lids during a 500ns simulation starting from the OO EM state (**a**, blue curves). The r.m.s.d. values of both RagB subunits decrease from ~30 Å in the EM structure (t = 0 ns) to ~15 Å, which is similar to the opening observed in one of the apo-RagAB simulations starting from the closed structure. Thus, it appears that the energy minimum for the open state in the simulations is different from that in solution, for reasons that are not clear.

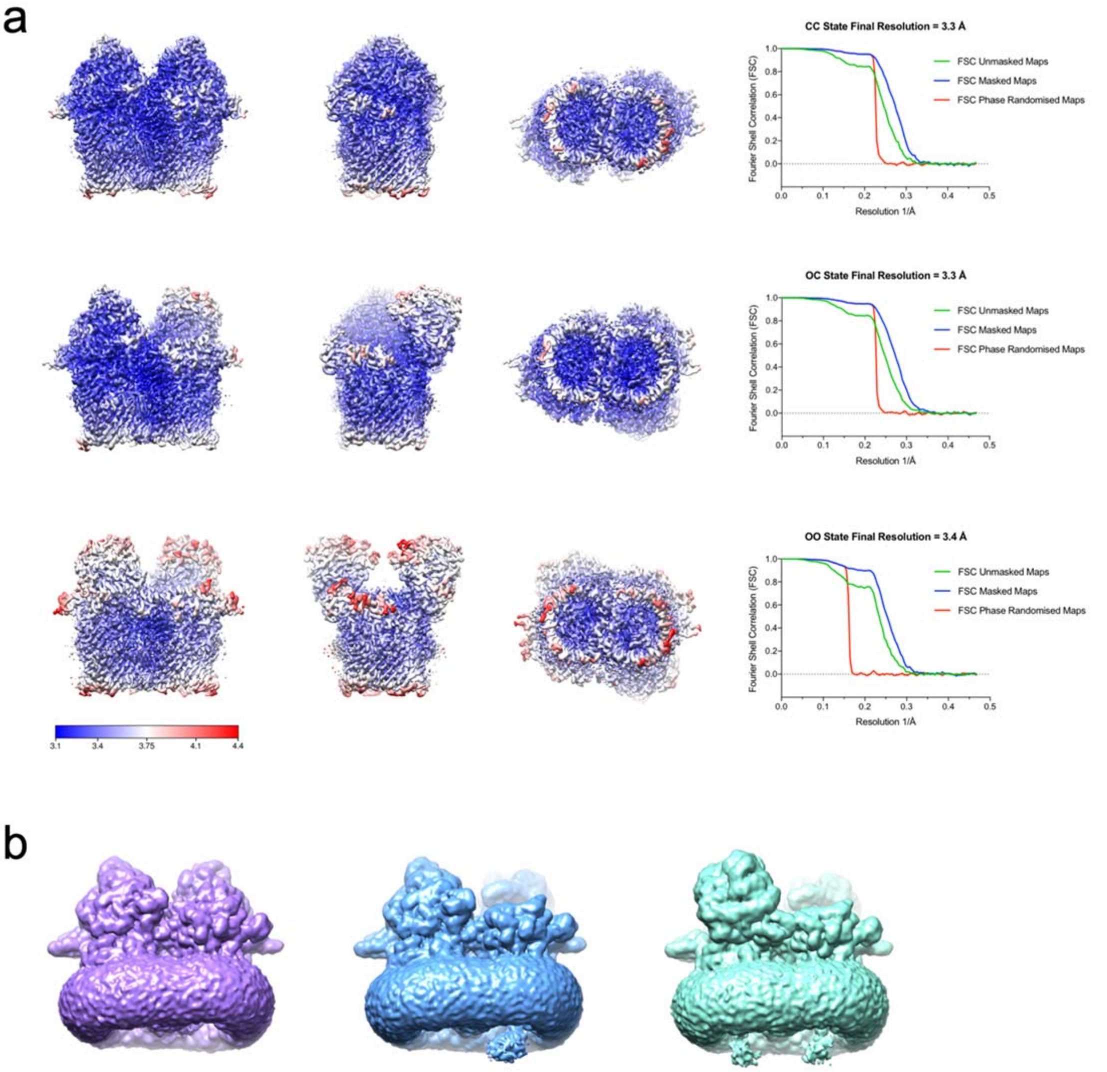


**Supplementary Figure 3 Local resolution-filtered cryoEM maps and evidence for DUF domain density.** **a**, CC, OC and OO states of RagAB filtered and coloured by local resolution. Corresponding FSC curves are shown (right). **b**, Unsharpened maps of RagAB displayed at low contour levels to reveal diffuse density attributed to the DUF domain. CC, OC and OO states are coloured purple, blue and green respectively.

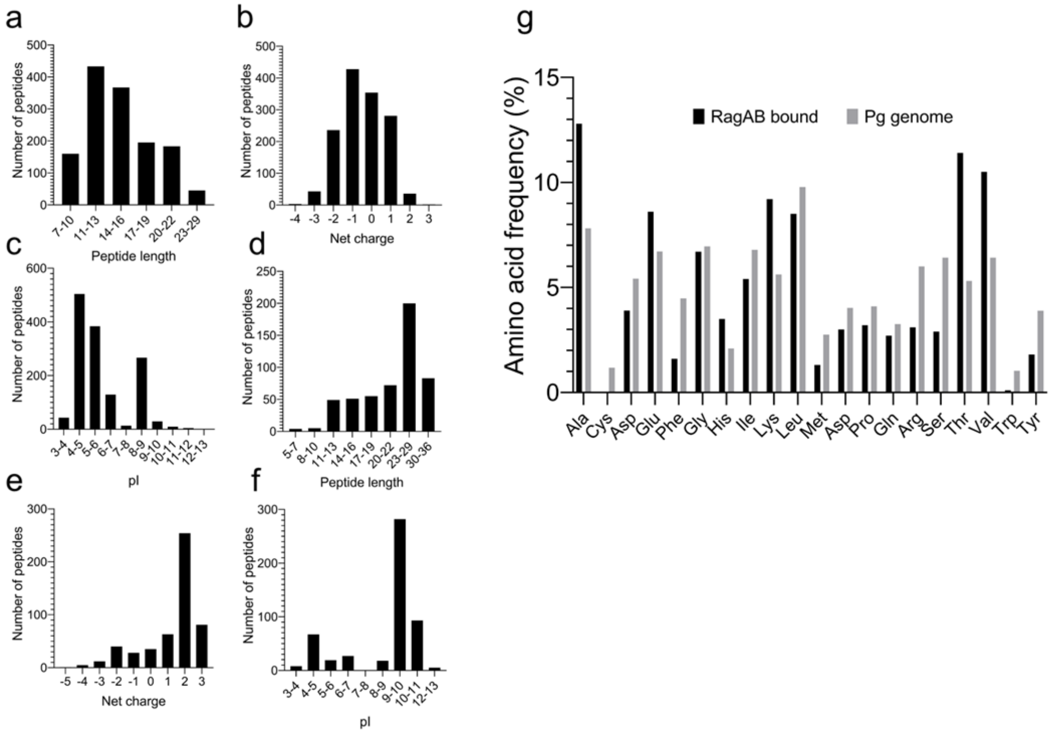


**Supplementary Figure 4 Semi-quantitative peptidomics of RagAB. a-c**, LC-MS/MS analysis of peptides bound to RagAB W83 KRAB, showing length distribution (**a**), total charge (**b**) and pI (**c**). **d-f** Analysis of peptides bound to RagAB W83 wild-type, showing length distribution (**d**), total charge (**e**) and pI (**f**). **g**, Amino acid frequency of RagAB-bound peptides (KRAB and wild-type combined; black) vs. the amino acid composition in the *P. gingivalis* proteome (gray).

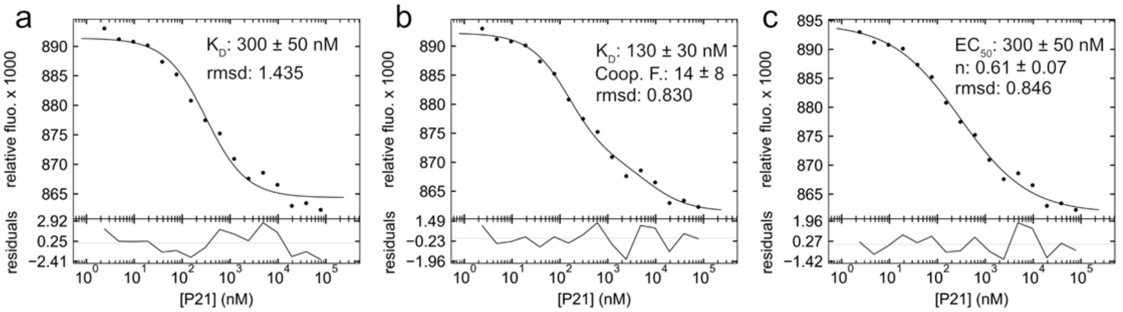


**Supplementary Figure 5 RagAB W83 exhibits negative cooperativity in binding of P21 peptide**. MST profiles for unlabelled P21 binding to His-tag labelled RagAB W83 with fitted line from 1:1 model (**a**), 1:2 Macro model (**b**) and Hill model (**c**). The residuals between the data and the fit line are indicated in the bottom panel. Experiments and listed $K_d$ and $EC_{50}$ values represent the mean of three independent experiments ± SD. Cooperativity factor (Coop. F.) and n factor were used to determine the mode of cooperativity. Root mean square deviation (rmsd) of the data from the fit line is indicated. Data were analysed in PALMIST version 1.4.0[5]. MST figures were rendered using GUSSI[6].

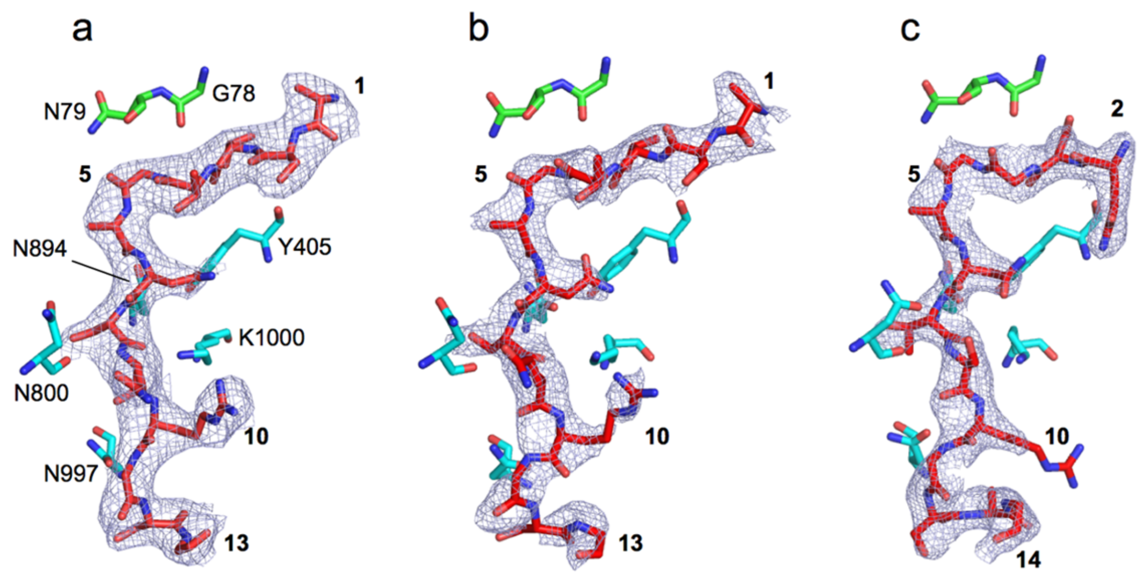


**Supplementary Figure 6 Electron density comparison for different peptide ensembles.** **a-c**, 2Fo-Fc electron density maps (contoured at 1.0 σ) for the peptide ensemble (red sticks) following final refinement for RagAB purified from W83 KRAB (**a**; 3.4 Å resolution), RagAB from W83 wild type (**b**; 3.0 Å) and RagAB from W83 wild type co-crystallised with excess P21 peptide (**c**; 2.6 Å). RagA and RagB residues that form hydrogen bonds with the backbone of the peptide are shown as cyan and green stick models, respectively. Peptide sequence of the P21 co-crystal structure was modelled as QNGGANTSRGSAG.

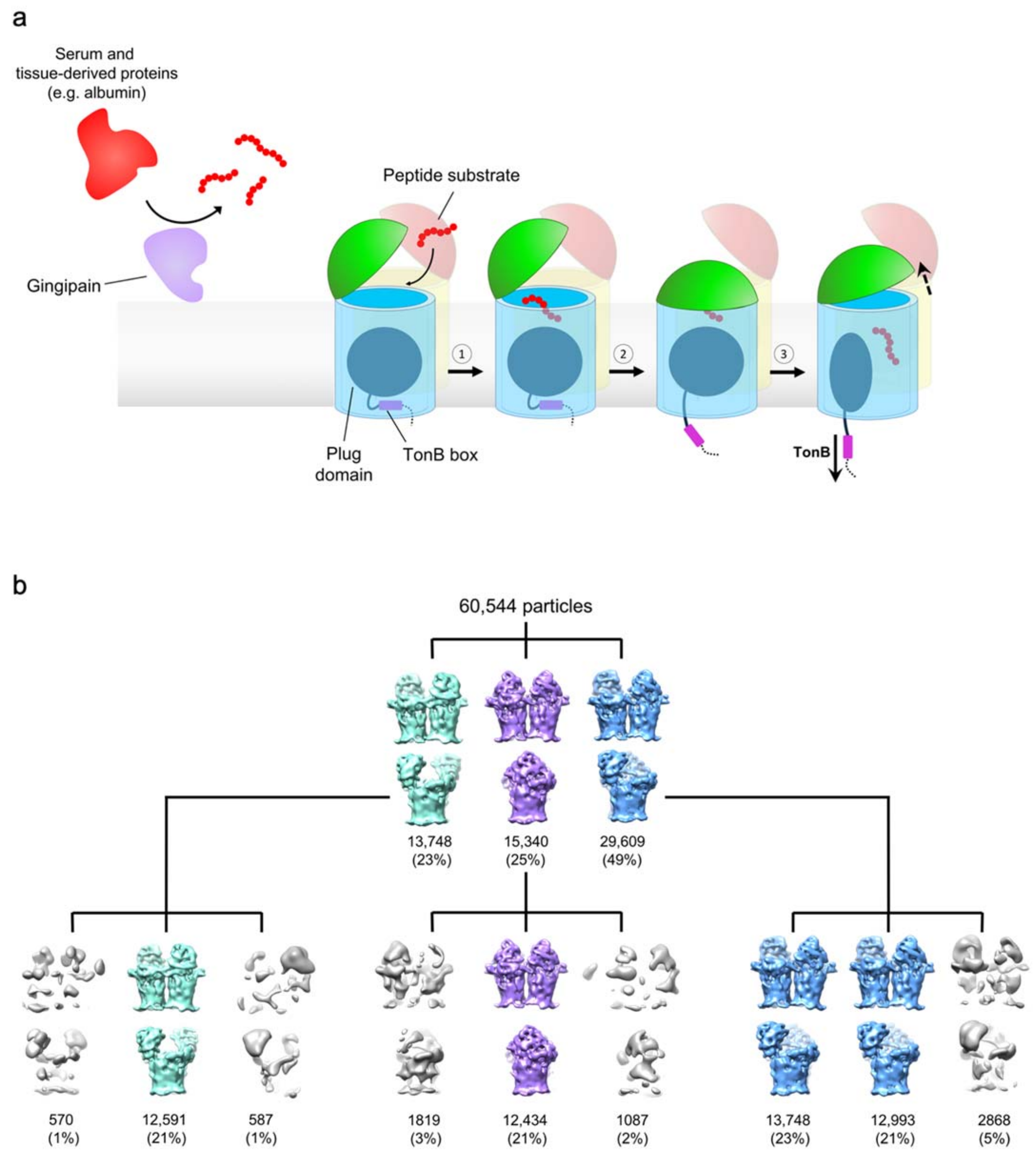
a
Serum and
tissue-derived proteins
(e.g. albumin)
Gingipain
Peptide substrate
Plug
domain
TonB box
TonB
b
60,544 particles
13,748
(23%)
15,340
(25%)
29,609
(49%)
570
(1%)
12,591
(21%)
587
(1%)
1819
(3%)
12,434
(21%)
1087
(2%)
13,748
(23%)
12,993
(21%)
2868
(5%)

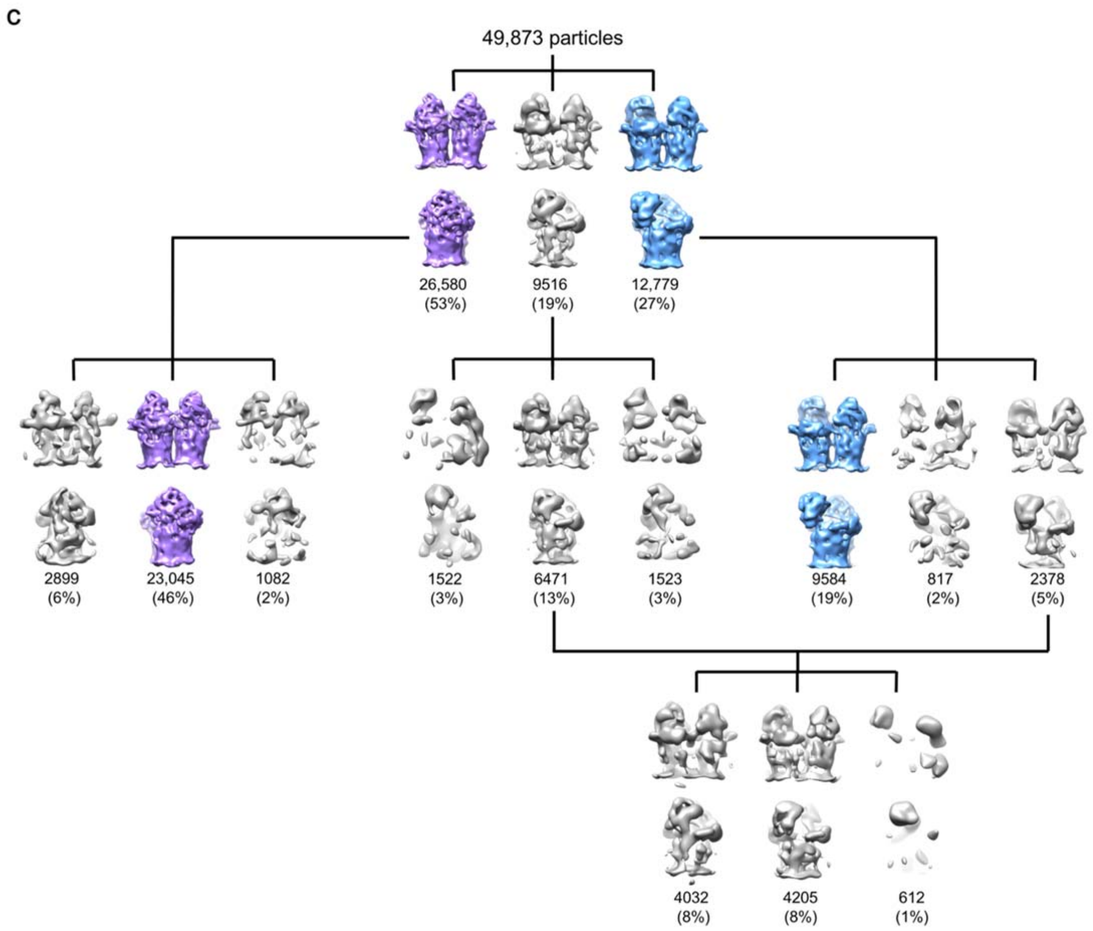


**Supplementary Figure 7 Schematic demonstrating the proposed mechanism of substrate capture and translocation by RagAB with supporting cryo-electron microscopy data. a**, Peptide ligands to be imported by the RagAB system are predominantly generated by the action of gingipains on serum and tissue-derived proteins. **1.** A lid-open state of RagAB permits peptide binding. **2.** Contributions from both RagA and RagB to peptide binding elicits closure of the lid, forming the transport-competent state of the complex. This is signalled across the OM by perturbation of the TonB box region on the periplasmic side of the plug domain, making it accessible to TonB. **3.** According to the literature consensus, TonB-mediated disruption of the plug permits substrate translocation and a return to the open state of RagAB. **b,c**, 3D classes for RagAB 'as purified' (**b**) and in the presence of 50-fold excess P21 peptide (**c**). Classes corresponding to the CC, OC and OO states are coloured purple, blue and green respectively. Junk or ambiguous classes *e.g.* where RagA barrels are incomplete are coloured grey. In the presence of P21 there was no clear OO state whilst the the proportion of the CC state increased, supporting the proposed mechanism of substrate capture.

**Supplementary Movie** Dynamics of the RagAB transporter. Cryo-EM map of open-closed (OC) RagAB, with cartoon models shown for RagA (blue) and RagB (green) subunits.

**Supplementary Table 1.** Crystallographic data collection and refinement statistics for RagAB.

| **Data collection** | W83-KRAB | W83-wild type | W83-wild type + P21 |
|---|---|---|---|
| DLS beamline/wavelength | i04-1/0.928 | i04/0.979 | i04-1/0.916 |
| Space Group | $C222_1$ | $P2_12_12_1$ | $P2_12_12_1$ |
| Cell dimensions | | | |
| a,b,c (Å) | 191, 377, 369 | 131, 142, 242 | 130, 143, 250 |
| □α,β,χ (°) | 90, 90, 90 | 90, 90, 90 | 90, 90, 90 |
| Molecules/AU (RagAB) | 4 | 2 | 2 |
| Resolution (Å) | 83.9-3.38 (3.44-3.38)[#] | 80.5-3.04 (3.09-3.04) | 142.7-2.61 (2.65-2.61) |
| I/ σI | 8.5 (2.0) | 3.7 (1.3) | 5.6 (1.0) |
| Completeness (%) | 99.6 (100) | 100 (99.5) | 99.9 (96.8) |
| Redundancy | 7.6 (8.1) | 7.4 (7.6) | 7.3 (7.6) |
| Rpim(%) | 8.4 (44.9) | 15.7 (55.8) | 10.2 (75.2) |
| CC (1/2) | 0.99 (0.74) | 0.95 (0.67) | 1.00 (0.53) |
| **Refinement** | | | |
| Resolution | 83.9-3.38 | 71.0-3.04 | 124.0-2.61 |
| $R_{work}/R_{free}$ (%) | 20.5/25.5 | 22.3/29.3 | 20.7/26.4 |
| Reflections | 184138 | 86882 | 140898 |
| No. Atoms | | | |
| Protein (RagA/RagB)* | 7048/3839 | 7069/3838 | 7064/3834 |
| Peptide* | 82 | 82 | 81 |
| B-factors (Å$^2$) | | | |
| Protein (RagA/RagB)* | 89/66 | 44/37 | 57/50 |
| Peptide* | 95 | 72 | 72 |
| Rmsd values | | | |
| Bond lengths (Å) | 0.011 | 0.011 | 0.008 |
| Bond Angles (°) | 1.37 | 1.33 | 1.03 |
| Molprobity clashscore | 15.1 | 14.4 | 9.7 |
| Ramachandran plot | | | |
| Favoured (%) | 86.4 | 92.0 | 93.8 |
| Outliers (%) | 2.4 | 0.6 | 0.6 |
| PDB code | 6SLI | 6SLJ | 6SLN |

[#] Parentheses are for the highest resolution shell
* Values are for the RagAB complex with the lowest B-factors

**Supplementary Table 2** CryoEM data acquisition parameters and model refinement statistics for RagAB

| | RagAB (principle dataset) | | | RagAB (control) | | | RagAB with P21 | | |
|---|---|---|---|---|---|---|---|---|---|
| **Data collection parameters** | | | | | | | | | |
| Detector | K2 Summit | | | K2 Summit | | | K2 Summit | | |
| Accelerating voltage (kV) | 300 | | | 300 | | | 300 | | |
| Magnification | 130,000 x | | | 130,000 x | | | 130,000 x | | |
| Total electron dose (e–/Å$^2$) | 77.88 | | | 75.1 | | | 75.3 | | |
| Number of frames | 48 | | | 50 | | | 50 | | |
| Dose per frame (e–/Å$^2$) | 1.62 | | | 1.5 | | | 1.5 | | |
| Pixel size (Å) | 1.07 | | | 1.07 | | | 1.07 | | |
| Defocus range (µm) | -1.2 to -3.0 | | | -1.5 to -3.3 | | | -1.5 to -3.3 | | |
| Micrograph movies | 3608 | | | 1212 | | | 1100 | | |
| **Image processing** | | | | | | | | | |
| Initial particle images | 491,870 | | | 74,857 | | | 82,062 | | |
| Conformation | **CC** | **OC** | **OO** | **CC** | **OC** | **OO** | **CC** | **OC** | **OO** |
| Final particle images | 86,877 | 213,143 | 51,849 | 12,434 | 26,741 | 12,591 | 23,045 | 9,584 | n/a |
| Applied symmetry | C2 | C1 | C2 | - | - | - | - | - | - |
| Map sharpening *B* factor (Å$^2$) | CC | OC | OO | - | - | - | - | - | - |
| Map resolution (Å) (FSC=0.143) | 3.3 | 3.3 | 3.4 | - | - | - | - | - | - |
| **Model refinement** | | | | | | | | | |
| PDB ID | 6SM3 | 6SMQ | 6SML | | | | | | |

| | | | |
|---|---|---|---|
| RMSD | | | |
| *Bond lengths (Å)* | 0.009 | 0.011 | 0.009 |
| *Bond angles (°)* | 0.938 | 0.998 | 0.996 |
| Validation | | | |
| *All-atom clashscore* | 2.45 | 3.32 | 3.67 |
| *MolProbity score* | 1.50 | 1.59 | 1.59 |
| *Rotamer outliers (%)* | 0.26 | 0.34 | 0.09 |
| Ramachandran plot | | | |
| *Favoured (%)* | 92.39 | 92.44 | 93.27 |
| *Allowed (%)* | 7.61 | 2.38 | 6.66 |
| *Outliers (%)* | 0 | 0.18 | 0.07 |
| Model vs map correlation | | | |
| *Cross correlation (mask)* | 0.87 | 0.87 | 0.83 |
| *Cross correlation (volume)* | 0.80 | 0.84 | 0.77 |

**Supplementary Table 3** RagAB peptidomics and *in vitro* binding of peptides to RagAB and RagB. Both analyses contain two spreadsheets: all information retrieved from Mascot (Mascot) and a reduced spreadsheet with summed spectra (duplicates) and one charge variant of each peptide (Spectral C.) For spectral C., additional statistics were calculated: Spectral count peptide - summed number of spectra of particular peptide; Spectral count protein - summed number of spectra per particular protein; Spectral count sample - summed number of spectra per particular sample; Ratio peptide/protein - ratio of total number of particular peptide spectra to total number of spectra per protein; Ratio peptide/sample - ratio of total number of particular peptide spectra to total number of spectra per protein.

**Supplementary Table 4** Primers, plasmids and strains used in this study.

#### Primers

| **Name** | **Sequence (5'→3')** |
| --- | --- |
| **RagBall plasmid** | |
| RagB_A_KpnI_F | ATTGGTACCATGAAAAAAATAATTTATTGGGTTG |
| RagB_A_BamHI_R | ATAGGATCCTTATATCGGCCAGTTCTTTATTAAC |
| RagB_tet_BamHI_F | ATAGGATCCACAACGAATTATCTCCTTAACGTACG |
| RagB_tet_XbaI_R | TCGTCTAGATTTTATTGCCAAGTTCTAATGCTTCT |
| RagB_B_XbaI_F | TGCTCTAGATTTAGTTGTAGATCTTACTATGAAA |
| RagB_B_SphI_R | CATGCATGCACAAAGATAAGATATCTGCC |
| **RagAall plasmid** | |
| RagA_A_SmaI_F | GTACCCGGGTGAAAAAAAGGATAATAGGATTAGTCT |
| RagA_A_NdeI_R | TATCATATGTTAGAAAGACAACTGAATACCCGC |
| RagA_erm_NdeI_F | TAACATATGATAGCTTCCGCTATTGCTTTTTTG |
| RagA_erm_XhoI_R | ATCCTCGAGTCTAGAGGATCCCCGAAGCTG |
| RagA_B_XhoI_F | AGACTCGAGGATTTACTTATTCTTAAGAAACATTTGATATGAA |
| RagA_B_SalI_R | CAGGTCGACGAAAGGGACTGCGCTCGG |
| **RagB-8His plasmid** | |
| RagB_8H_Fl | CACCATCACCATCACCATCACCATTAAGGATCCACAACGAATTATCTC |
| RagB_8H_Fs | TAAGGATCCACAACGAATTATCTC |
| RagB_8H_Rl | ATGGTGATGGTGATGGTGATGGTGTATCGGCCAGTTCTTTATTAACTG |
| RagB_8H_Rs | TATCGGCCAGTTCTTTATTAACTG |
| **Δ*ragB* plasmid** | |
| delRagB_KpnI_F | CAGTGGTACCCCAATTCGTTCTATATGGCT |
| delRagB_BamHI_R | GTTGTGGATCCATCAAATGTTTCTTAAGAATAAGTAAATC |
| **Δ*ragA* plasmid** | |
| delRagA_A_F | TGAATTCGAGCTCGGTACCCGACACGAAGGAGTTTATTGCG |
| delRagA_A_R | AAAGCAATAGCGGAAGCTATTCTAAGCAATTTGCTCACCATAC |
| delRagA_erm_F | TGGTGAGCAAATTGCTTAGAATAGCTTCCGCTATTGCTTTTTT |
| delRagA_erm_R | TTCTTAAGAATAAGTAAATCTCTAGAGGATCCCCGAAGCT |
| delRagA_B_F | GCTTCGGGGATCCTCTAGAGATTTACTTATTCTTAAGAAACATTTGAT |
| delRagA_B_R | GTCGACTCTAGAGGATCCCCTTTGTCGATCTCGCTGTG |
| **Δ*ragAB* plasmid** | |
| delRagAB_KpnI_F | CAGTGGTACCGACACGAAGGAGTTTATTGCG |
| delRagAB_BamHI_R | GTTGTGGATCCTCTAAGCAATTTGCTCACCATAC |
| **RagB$_{BL}$ plasmid** | |
| RagB_DEDEloop_Fl | GGACGTGCTCGTAAGCGTAAGGTCTCCGGTATAGCTGGCTACTATT |
| RagB_DEDEloop_Rl | GGAGACCTTACGCTTACGAGCACGTCCATGGTCGTTA |
| RagB_DEDEloop_Fs | GTCTCCGGTATAGCTGGCTACTATT |
| RagB_DEDEloop_Rs | AGCACGTCCATGGTCGTTA |
| **$\Delta_{AL}$ plasmid** | |
| RagB_delAcLoop_Fl | GGCATACTTAACGACCATGGAGGTATAGCTGGCTACTATTTCGTAT |
| RagB_delAcLoop_Rl | GAAATAGTAGCCAGCTATACCTCCATGGTCGTTAAGTATGC |
| RagB_delAcLoop_Fs | GGTATAGCTGGCTACTATTTCGTAT |
| RagB_delAcLoop_Rs | TCCATGGTCGTTAAGTATGC |
| **RagAB$_{mono}$ plasmid** | |
| RagAmono6H_Fl | GCCAGCATCATCATCATCATCATGGAAAAACCGGAAATAGTTTG |
| RagAmono6H_Rl | TTCCATGATGATGATGATGATGCTGGCTCAGTAACATCAACTTATC |
| RagAmono6H_Fs | GGAAAAACCGGAAATAGTTTG |
| RagAmono6H_Rs | CTGGCTCAGTAACATCAACTTATC |
| **Δhinge1 plasmid** | |
| RagBdelHinge1_Fl | GAGATTGGTAATTACAACCACAATCCCGACCTCTCGTGG |
| RagBdelHinge1_Rl | TCCCACGAGAGGTCGGGATTGTGGTTGTAATTACCAATCTCCGAG |
| RagBdelHinge1_Fs | AATCCCGACCTCTCGTGG |
| RagBdelHinge1_Rs | GTGGTTGTAATTACCAATCTCCGAG |
| **Δhinge2 plasmid** | |
| RagBdelHinge2_Fl | CGCACTACGAATGATATGGGCGTAGGCTCTATGAAAAATACGGG |
| RagBdelHinge2_Rl | ATTTTTCATAGAGCCTACGCCCATATCATTCGTAGTGCGGAC |
| RagBdelHinge2_Fs | GTAGGCTCTATGAAAAATACGGG |
| RagBdelHinge2_Rs | CATATCATTCGTAGTGCGGAC |
| **ΔTonB plasmid** | |
| RagAdelTonB_Fl | GTACTGGATCCGGACTCTAAGGGTACGGGACAGAAACTCAG |
| RagAdelTonB_Rl | GCTGAGTTTCTGTCCCGTACCCTTAGAGTCCGGATCCAGTACG |
| RagAdelTonB_Fs | GGTACGGGACAGAAACTCAG |
| RagAdelTonB_Rs | CCTTAGAGTCCGGATCCAGTACG |

| | |
|---|---|
| **RagB W83 in ATCC plasmid** | |
| ATCC_RagB_vect_F | ACAACGAATTATCTCCTTAACG |
| ATCC_RagB_vect_R | AATGATTTACTGTTAATCGTTAGTAC |
| RagBW83inATCC_F | ACGATTAACAGTAAATCATTATGAAAAAAATAATTTATTGGGTTGC |
| RagBW83inATCC_R | TAAGGAGATAATTCGTTGTTTATATCGGCCAGTTCTTTATTAAC |
| **Δ*ragB*-ATCC plasmid** | |
| delRagB_ATCC_A_F | TGAATTCGAGCTCGGTACCCTTTTTGGAACAATCGTTTG |
| delRagB_ATCC_A_R | TTAAGGAGATAATTCGTTGTAATGATTTACTGTTAATCGTTAGTACTC |
| delRagB_ATCC_tet_F | ACGATTAACAGTAAATCATTACAACGAATTATCTCCTTAACGTACG |
| delRagB_ATCC_tet_R | CGTCGAAACATCTAATTGAATTTTATTGCCAAGTTCTAATGCTTC |
| delRagB_ATCC_B_F | ATTAGAACTTGGCAATAAAATTCAATTAGATGTTTCGACG |
| delRagB_ATCC_B_R | GTCGACTCTAGAGGATCCCCTACGGAGAACATATGTTCC |
| **Δ*ragAB*-ATCC plasmid** | |
| delRagAB_ATCC_A_F | TGAATTCGAGCTCGGTACCCTTTCCGTCTCTCTTATGACGAAGAG |
| delRagAB_A_R | TTAAGGAGATAATTCGTTGTTCTAAGCAATTTGCTCACCATAC |
| delRagAB_ATCC_tet_F | TGGTGAGCAAATTGCTTAGA ACAACGAATTATCTCCTTAACGTAC |
| delRagAB_ATCC_tet_ | CGTCGAAACATCTAATTGAATTTTATTGCCAAGTTCTAATGCTTC |
| delRagAB_ATCC_B_F | ATTAGAACTTGGCAATAAAATTCAATTAGATGTTTCGACG |
| delRagAB_ATCC_B_R | GTCGACTCTAGAGGATCCCCTACGGAGAACATATGTTCC |
| **RagAB-W83-pTIO plasmid** | |
| RagAB_W_pTIO_F | GCAGCCCGGGTTGGCGCGAGAAGTAAAAAAATC |
| RagAB_W_pTIO_R | CCGCTCTAGATTATATCGGCCAGTTCTTTATTAACTG |
| **ΔDUF plasmid** | |
| RagAdelDUF_Fl | GCTATGGCCCAGAATAGAACCGTTCTGGAGCAGGTAGTTGTAT |
| RagAdelDUF_Rl | TACAACTACCTGCTCCAGAACGGTTCTATTCTGGGCCATAG |
| RagAdelDUF_Fs | GTTCTGGAGCAGGTAGTTGTAT |
| RagAdelDUF_Rs | ACGGTTCTATTCTGGGCCATAG |

### Plasmids

| Plasmid | Relevant features | Source |
|---|---|---|
| **pUC19** | *E. coli* cloning vector, $Ap^R$ | Thermo Scientific |
| **pTIO-1** | *E. coli-Bacteroides* shuttle vector, $Ap^R$ | [7] |
| **RagBall** | Master plasmid for RagB modifications, derivative of pUC19 | This study |
| **RagB-8His** | Plasmid for insertion of 8xHis at C-terminal of RagB, used for purification of RagAB complex, derivative of RagBall, | This study |
| **Δ*ragB*** | Plasmid for deletion of *ragB* gene, derivative of RagBall | This study |
| **Δ*ragA*** | Plasmid for deletion of *ragA* gene, derivative of RagAall | This study |
| **Δ*ragAB*** | Plasmid for deletion of *ragA* and *ragB* genes, derivative of delRagB | This study |
| **RagAall** | Master plasmid for RagA modifications, derivative of pUC19 | This study |
| **$RagB_{BL}$** | Plasmid for substitution of $D^{99}$-$E^{102}$ with RKRK in *ragB* gene, derivative of RagBall | This study |
| **$\Delta_{AL}$** | Plasmid for deletion of acidic loop $R^{97}$-$S^{104}$ in *ragB* gene, derivative of RagBall | This study |
| **$RagAB_{mono}$** | Plasmid for insertion of 6xHis after $Q^{570}$ in *ragA* gene, derivative of RagAall | This study |
| **Δhinge1** | Plasmid for deletion of $Q^{670}$- $G^{691}$ in *ragA* gene, derivative of RagAall | This study |
| **Δhinge2** | Plasmid for deletion of $L^{731}$ -$N^{748}$ and insertion of glycine in postion 731 in *ragA* gene, derivative of RagAall | This study |
| **ΔTonB** | Plasmid for deletion of $V^{100}$-$Y^{109}$ in *ragA* gene, derivative of RagAall | This study |
| **ΔDUF** | Plasmid for deletion of $V^{25}$-$K^{99}$ in *ragA* gene, derivative of RagAall | This study |
| **RagB W83 in ATCC** | Plasmid for substitution of *ragB* from ATCC33277 with *ragB* from W83 strain | This study |
| **Δ*ragB*-ATCC** | Plasmid for deletion of *ragB* gene in ATCC33277 strain | This study |
| **Δ*ragAB*-ATCC** | Plasmid for deletion of *ragA* and *ragB* genes in ATCC33277 strain | This study |
| **RagAB-W83-pTIO** | Shuttle plasmid for expression of *ragAB* genes from W83 strain | This study |

### Strains

| Strain | Relevant genotype | Source |
|---|---|---|
| | ***E. coli*** | |
| **DH5α** | *fhuA2* Δ(*arg*F-*lac*Z)U169 *pho*A *gln*V44 Φ80 Δ(*lac*Z)M15 *gyr*A96 *rec*A1 *rel*A1 *end*A1 *thi*-1 *hsd*R17 | New England Biolabs |
| **BL21 (DE3)** | *fhuA2 [lon] ompT gal (λ DE3) [dcm] ΔhsdS λ DE3 = λ sBamHIo ΔEcoRI-B int::(lacI::PlacUV5::T7 gene1) i21 Δnin5* | Invitrogen |
| **S-17 λpir** | *creC510 hsdR17 thiE1 endA1 recA1 LAMpir pro-82 RP4-2(Km::Tn7,Tc::Mu-1)* | [8] |
| | ***P. gingivalis*** | |
| **W83** | Wild type | [9] |
| **ATCC33277** | Wild type | [10] |
| **HG66** | Wild type | [11] |
| **A7436** | Wild type | [12] |
| **381** | Wild type | [13] |
| **KRAB** | *rgpA rgpB kgp* ($Cm^R$)($Em^R$)($Tet^R$) | [14] |
| ***ΔragA*** | *ragA* (NCBI:PG_0185)($Em^R$) | This study |
| **Δ*ragB*** | *ragB* (NCBI:PG_0186)($Tet^R$) | This study |
| **Δ*ragAB*** | *ragA ragB* ($Tet^R$) | This study |
| **ΔTonB** | *ragA* $\Delta^{101-108}$ ($Em^R$) | This study |
| **$RagAB_{mono}$** | *ragA* p.$Q^{570}$_insHHHHHH_$G^{571}$ ($Em^R$) | This study |
| **Δhinge1** | *ragA* $\Delta^{670-691}$($Em^R$) | This study |
| **$\Delta_{AL}$** | *ragB* $\Delta^{97-104}$($Tet^R$) | This study |
| **$RagB_{BL}$** | *ragB*p.D99R;E100K;D101R;E102K($Tet^R$) | This study |
| **ΔDUF** | *ragA* $\Delta^{25-99}$ ($Em^R$) | This study |
| **Δhinge2** | *ragA* $\Delta^{670-691}$($Em^R$) | This study |
| **RagB-8His** | *ragA*p.L731G;$\Delta^{731-748}$;($Em^R$) | This study |
| **RagB W83 in ATCC 33277** | *ragB*(NCBI:PGN_0294):*ragB*(NCBI:PG_0186)($Tet^R$) | This study |
| ∆***ragAB* ATCC 33277** | *ragA*(NCBI: PGN_0293*); ragB*(NCBI: PGN_0294)($Tet^R$) | This study |
| **RagAB W83 in ATCC 33277** | *ragA*(NCBI: PGN_0293*); ragB*(NCBI: PGN_0294)($Tet^R$)/*ragA* (NCBI:PG_0185) *ragB* (NCBI:PG_0186) ($Em^R$) | This study |
